# On the mode of anti-*C. albicans* activity of a bis(benzoxaborole) analogue of Tavaborole

**DOI:** 10.1101/2024.07.29.605614

**Authors:** Sachin Gadakh, Teresa Szczepińska, Małgorzata Potoczna, Shiho Okitsu-Sakurayama, Piotr Podlasz, Marta Rogalska, Ewa Kaczorowska, Agnieszka Adamczyk-Woźniak, Monika Staniszewska

## Abstract

We presented the pharmacodynamic relationship between benzoxaborole concentrations and *Candida albians* blastoconidial population dynamics. Bis(benzoxaborole) analogue (**2**) and **Tavaborole** (ref.) showed comparable moderate effects against *C. albicans* (time-kill kinetic assays). Benzoxaboroles inhibited the *C. albicans* growth during 72 h (fungicidal after 2 h of that time) with metabolic reduction (%R=64). Biofilm-inhibiting concentration (BIC_50_=2 μg/mL) is the lowest concentration of bezoxaboroles that presented 50% inhibition of biofilm metabolic activity vs non-treated control. **2** displayed poor ability to inhibit morphogenesis of *C. albicans*. Safety and fungicidal activity are still in high demand against biofilm grown on fibroblasts and the zebrafish model *in vivo*. This biofilm model was used to study the interactions between the *C. albicans* morphogenesis and benzoxaboroles. Benzoxaboroles displayed selectivity in cytotoxic effects. **2** exhibited significantly lower embryotoxicity vs ref. IC_50_>128 μg/mL for **2**. Ref. at 256 μg/mL showed approximately 80% viability of VERO E6 cells. A higher selectivity of the ref—drug to the pathogen than to the mammalian cells was observed. Contrariwise, ref. and **2** showed IC_50_=2 μg/mL against PBMCs. Benzoxaborole antifungal development targets ergosterol binding. RNAseq data indicated that efflux pumps (MDR) in *C. albicans* were upregulated. Inositol-1-phosphate synthase was repressed under the benzoxaborole treatment. *MDR1* upregulation by **2** was accompanied by the *IFD6* (aldo-keto reductase) increase and the coordination of multiple coactivators (*IFD6, TNA1* encoding putative nicotinic acid transporter). Benzoxaboroles represent a similar resistance mechanism to azoles due to the subsequent expression of *MDR1* and *IDF6*. Docking studies confirmed the proposed interactions of benzoxaborole adenosinemonophosphate adduct with LeuRS. Moreover, *C5_04480C_A* (cell wall biogenesis, protein folding, modification, and destination) was negatively regulated in response to the benzoxaborole stress. Benzoxaborole-altering efflux inhibitors are important for the development of combination strategies in candidiasis. Our findings present an innovative concept that can inspire further studies for designing and building new antifungal benzoxaborole.

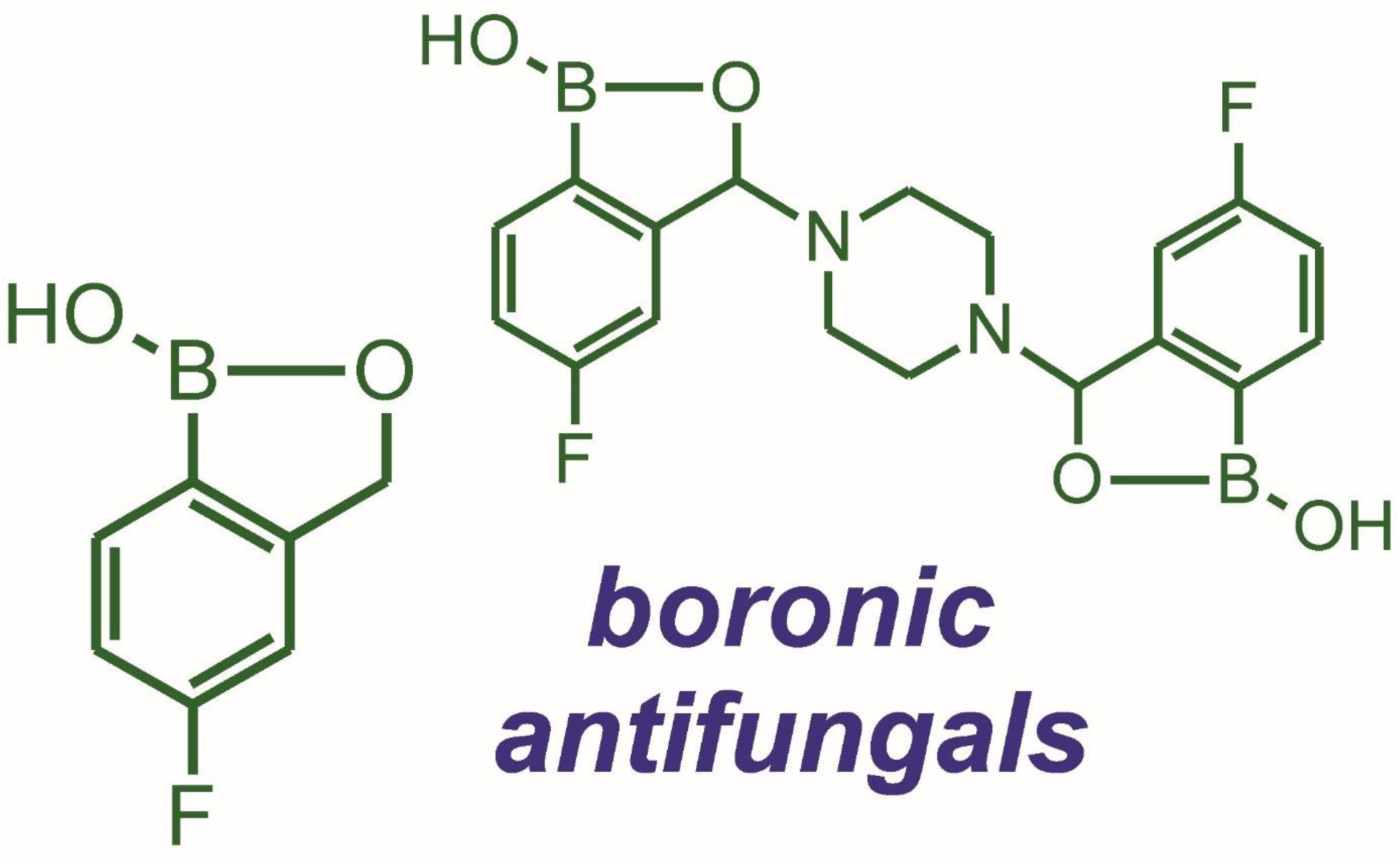

## Introduction

Epidemiological data obtained over the last 20 years show that strains of the genus *Candida spp.* are the main etiological factor of fungal infections occurring among hospitalized patients[1]. Fungal infections have a high mortality rate (from 30 to 50%) among immunocompromised patients (HIV, cancer, and transplant patients)[1],[2],[3]. In addition, ineffective antifungal therapy contributes to an 85% increase in the costs of medical care for a patient with candidemia[4]. Moreover, the increasing use of antimycotics currently used in medicine (e.g. echinocandins) is associated with the acquisition of resistance by *Candida* strains. At the same time, therapy with amphotericin B is limited by undesirable side effects, such as nephro[5]- and hepatotoxicity.[6] Therefore, searching for new compounds with high antifungal activity and safe for patients is a worthwhile task, both from the medical as well as socioeconomic points of view. Benzoxaboroles have gained great popularity in the last decade due to their antimicrobial activity.[7],[8],[9],[10] Several years of clinical research resulted in the approval of Tavaborole – the first known antifungal drug of a benzoxaborole structure.[11],[12],[13],[14],[15],[16] This compound is currently used in the topical treatment of fungal infections called onychomycosis. Several analogs of Tavaborole have been recently obtained and studied as potential antimicrobial agents.[17],[18],[19],[20],[21] Among them bis(benzoxaboroles) have been scarcely studied, yet showing a promising antimicrobial action.[19],[21],[22],[23] This work aimed to investigate the antifungal action mode of **2** and the Tavaborole reference drug (ref.). Both compounds were tested against the *C. albicans* hyphal growth at different stages of biofilm formation on the abiotic surface. While the integrity of fungal cells requires the presence of the membrane, the studies involving an ergosterol that provides mechanical support and protection against different environmental stresses [24] were included in the study. Thus, we examined whether the increased sensitivity of the *C. albicans* cells to benzoxaboroles was due to ergosterol being more accessible to the agent. We wanted to understand whether bezoxaboroles regulate sterol levels. Finally, we proved the benzoxaborole action mode in anti-virulence potential by screening the *C. albicans* calcineurin mutants. It is important to mention that calcineurin is required for the formation of invasive hyphae and pathogenicity.[25] Thus we used these genetic tools (*Δcnb1/Δcnb1* and *CNB1/Δcnb1*[25]) to prove the benzoxaborole action modes. Therefore, the study aimed to evaluate the impact of bezoxaboroles against the *C. albicans* wt and mutant biofilm generated on fibroblast. The overall effect of the *C. albicans* biofilm formation on fibroblasts under the benzoxaborole treatment has not yet been studied. Further knowledge about these interactions may help develop new therapies to control *C. albicans* pathogenesis. Using the high-throughput next-generation sequencing technique RNA-seq, we also uncovered the underlying virulence factors whose expressions are suppressed by benzoxaborole treatments and revealed key cell cycle proteins involved in the yeast-to-hyphal transition, as the potential antifungal drug targets. The cytotoxicity of **2** and ref. was evaluated against the epithelial cells VERO E6 and peripheral blood mononuclear cells (PBMCs). To the best of our knowledge, the cytotoxicity of the studied compounds towards PBMCs is being reported here for the first time.

The studied bis(benzoxaborole) (**2**) combines two Tavaborole units connected by a piperazine fragment. Its anti-*C. albicans* activity was not evaluated in detail previously, apart from promising yet preliminary reports on the agar diffusion method studies.[19] The bis(benzoxaborole) Tavaborole analogue (**2**) was obtained according to a modified solution method starting from 4-fluoro-2-formylphenylboronic acid (**1**).[26] The same starting material was used to obtain a Tavaborole sample for comparative studies (Scheme 1).[27]

**Scheme 1:**
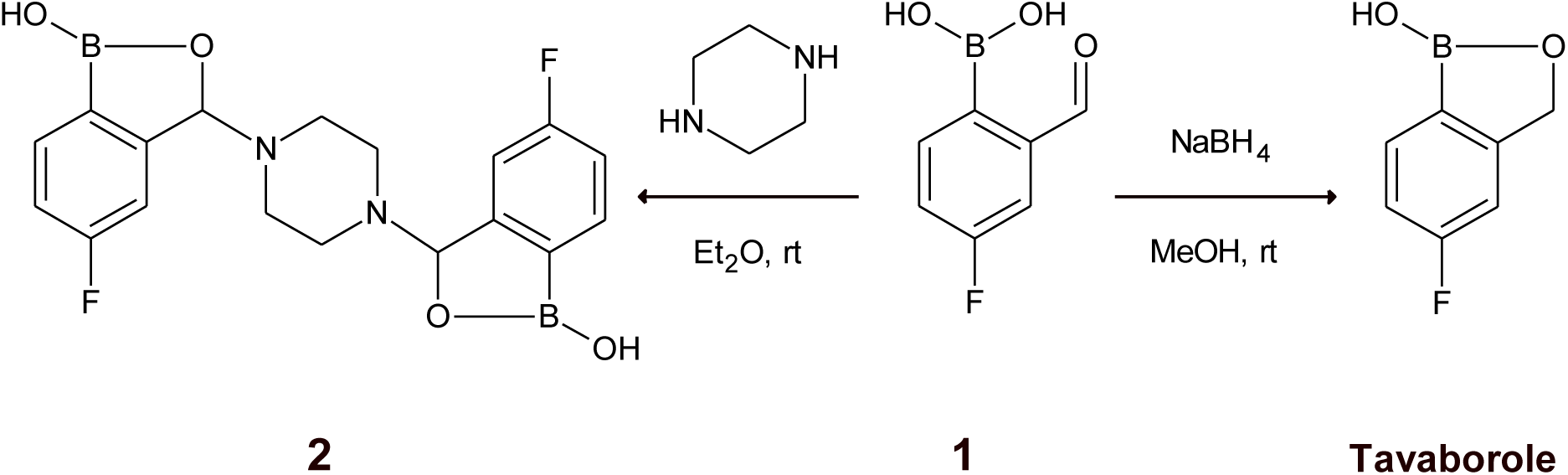
Synthesis of Tavaborole and bis(benzoxaborole) **2** starting form **1**.

### Synthesis of 2

Two solutions were prepared: a solution of 4-fluoro-2-formylphenylboronic acid (**1**) (9.01 g, 52.97 mmol) in diethyl ether (600 mL) and a solution of piperazine (2.28 g, 26.49 mmol) in diethyl ether (200 mL). Next, a solution of piperazine was added dropwise to a stirred solution of 4-fluoro-2-formylphenylboronic acid (**1**). After the addition of ca 80 mL of the piperazine solution, the first appearance of white precipitate was noticed. After the addition of the entire piperazine solution, stirring was turned off and the mixture was left in a closed flask for 24 hours at room temperature. The obtained solid was filtered, thoroughly washed with diethyl ether (20 mL) and dried in air for 3 days at room temperature. After this time the white, solid product was weighed and next analyzed (9.69 g, 24.47 mmol, yield: 92.4%).

**^1^H NMR** (500 MHz, DMSO-d_6_): δ (ppm) = 9.23 (s, 1H, B-OH), 9.20 (s, 1H, B-OH), 7.75-7.72 (dd, 1H, Ph), 7.71-7.68 (dd, 1H, Ph), 7.26-7.22 (ddd, 1H, Ph), 7.21-7.17 (ddd, 1H, Ph), 7.12-7.10 (dd, 1H, Ph), 7.08-7.06 (dd, 1H, Ph), 5.83 (s, 1H, CH), 5.79 (s, 1H, CH), 2.60 (br s, 4H, CH_2_CH_2_), 2.43 (br s, 4H, CH_2_CH_2_);**^19^F NMR** (470 MHz, DMSO-d_6_): δ (ppm) = -109.65, -109.78; **^11^B NMR** (160 MHz, DMSO-d_6_): δ (ppm) = 15

### Synthesis of Tavaborole

To the solution of 1.0965 g of the 2-formyl-4-fluorophenylboronic acid (**1**) in 13 mL of methanol, NaBH_4_ (0.2468g) was added portion-wise resulting in heating and bubbling. The reaction mixture was left for 24 h at room temperature. After that time a 3M HCl_aq_ was added (about 5 mL) resulting in a pH of about 2 (control with a universal pH indicator paper). Mixing was continued for about 20 min followed by removal of the volatiles at rotary evaporator. The resulting white precipitate was filtered off and washed thoroughly with distilled water till neutral reaction of the filtrate. The solid was dried on air at room temperature yielding 0.7543 g of pure Tavaborole (76% yield). The purity of the sample was confirmed based ^1^HNMR, ^19^F and ^11^B NMR, showing exclusively signals corresponding to the desired molecule as well as elemental analysis: Calculated for C_7_H_6_BFO_2_: C [%]: 55.34, H [%]: 3.98. determined: C [%]: 55.5, H [%]: 3.90.

**^1^H NMR** (500 MHz, (CD_3_)_2_CO) δ 8.12 (s, OH), 7.76 (dd, *J* = 8.1, 5.7 Hz, Ar), 7.24 – 7.15 (m, Ar), 7.15 – 7.07 (m, Ar), 5,02 (s, CH_2_); **^19^F NMR** (470 MHz, (CD_3_)_2_CO) δ – 111,87 (m); **^11^B NMR** (160 MHz, (CD_3_)_2_CO) δ 32 (s), mp. 124-130°C

### Activity against planktonic and sessile *C. albicans* morphotypes

The percentage of cell growth reduction (%R) determined for **2** and **Tavaborole** (ref) is shown in **Table 1**. 100 % R for **2** was determined at 64 µg/mL versus ref. at 16 µg/mL. In **Fig. 1A**., we evaluated the *C. albicans* metabolic activity (cell viability) inhibition to approximately 80% by **2** at 64 μg/mL versus 60% even at the lower concentration of ref. (16 μg/mL). The presence of exogenous ergosterol in the culture medium implies benzoxaboroles have the potential to target ergosterol (**Fig. 1B**). The 48-hour treatment of the *C. albicans* cells with benzoxaboroles in the presence of exogenous ergosterol resulted in cell viability in the range from 98.5%±0.03 to 100.00% (**Fig. 1B**).

**Fig. 1.**
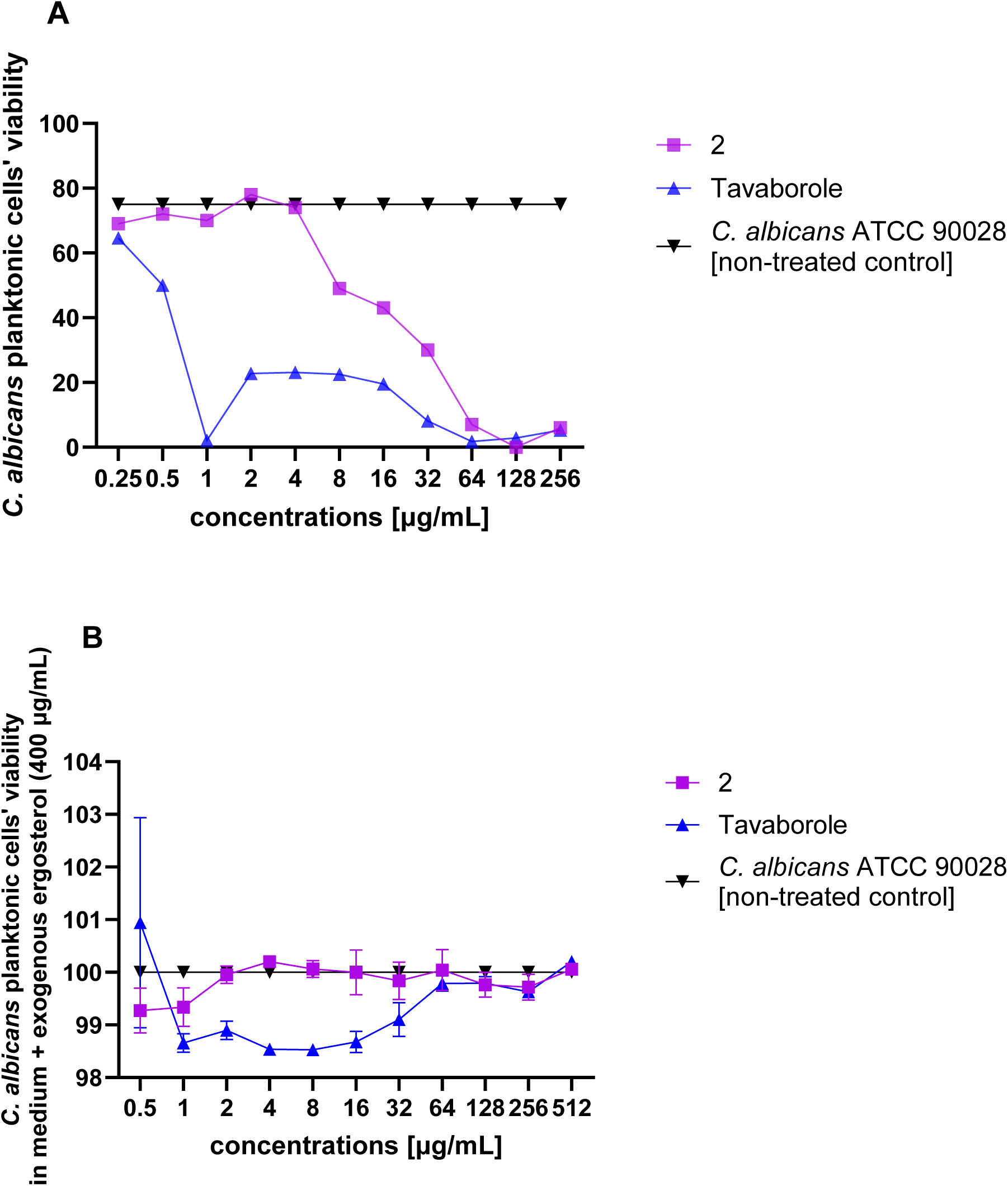
*Candida albicans* metabolic activity reduction by benzoxaborole derivatives Influence of bezoxaboroles concentrations on metabolic activity of blastoconidial cells. MTS test (3- (4.5-dimethylthiazol-2-yl)-5-(3-carboxymethoxyphenyl)-2-(4-sulfophenyl)-2H-tetrazolium (MTS, Promega, USA) was used to assess *C. albicans* proliferation. (**A**) The suspension of *C. albicans* cells in medium YEPD at the density of 1-3 x 10^2^ CFU/mL was incubated with compounds **2** and **Tavaborole** at the range of conc.: 0.25-256 μg/mL for 48 h at 37^°^C. (**B**) The activity of benzoxaboroles against *C. albicans* growing in a medium with exogenous ergosterol (exErg) at 400 µg/mL for 48 h. %viable cells = OD490 test sample/OD490 control x 100%, control means cells treated with 0.96% DMSO. The results from 3 independent experiments were analyzed. Statistically significant differences were noted between non-treated growth and treated samples (P< 0.05, Unpaired t-test).

**Table 1.** The cell growth inhibition (%R) of planktonic *C. albicans* ATCC 90028 *In vitro* determination of *Candida albicans* 90028 growth reduction for **2** and Tavaborole according to the American CLSI (Clinical and Laboratory Standards Institute[28]). *Blastoconidial optical density was reduced at the highest concentration tested of **2** (%R=100 at 64 µg/mL) vs Tavaborole (%R=100 at 16 µg/mL).

|  |  | **2** | **Tavaborole** |
| --- | --- | --- | --- |
| Conc. (µg/mL) | Incubation time (h) |  |  |
| 64 | 18 | 100.00 ± 0.00 | 100.00 ± 0.00 |
|  | 48 | 100.00 ± 0.00* | 100.00 ± 0.00 |
| 32 | 18 | 99.94 ± 0.10 | 100.00 ± 0.00 |
|  | 48 | 99.70 ± 0.00 | 100.00 ± 0.00 |
| 16 | 18 | 98.52 ± 0.93 | 100.00 ± 0.00 |
|  | 48 | 99.71 ± 0.16 | 100.00 ± 0.00* |
| 8 | 18 | 99.98 ± 0.00 | 99.88 ± 0.01 |
|  | 48 | 99.22 ± 0.00 | 99.99 ± 0.00 |
| 4 | 18 | 98.78 ± 1.86 | 99.08 ± 1.43 |
|  | 48 | 99.12 ± 0.04 | 99.99 ± 0.00 |
| 2 | 18 | 98.57 ± 1.73 | 98.59 ± 2.40 |
|  | 48 | 99.09 ± 0.03 | 99.99 ± 0.00 |
| 1 | 18 | 97.19 ± 2.32 | 97.66 ± 2.03 |
|  | 48 | 99.06 ± 0.01 | 99.87 ± 0.04 |
| 0.5 | 18 | 97.87 ± 2.21 | 97.34 ± 2.31 |
|  | 48 | 99.06 ± 0.03 | 99.39 ± 0.03 |

In **Fig. 2**, the *C. albicans* pattern of growth and kill by **2** and **Tavaborole** (ref.) is shown. The time-kill curves showed a log phase of 6–18 h and their maximum growth was lower than the control growth for *C. albicans*. In general, growth curves for *C. albicans* showed an increase in blastoconidial population size compared to that observed for *C. albicans* when exposed to **2** or ref.

**Fig. 2.**
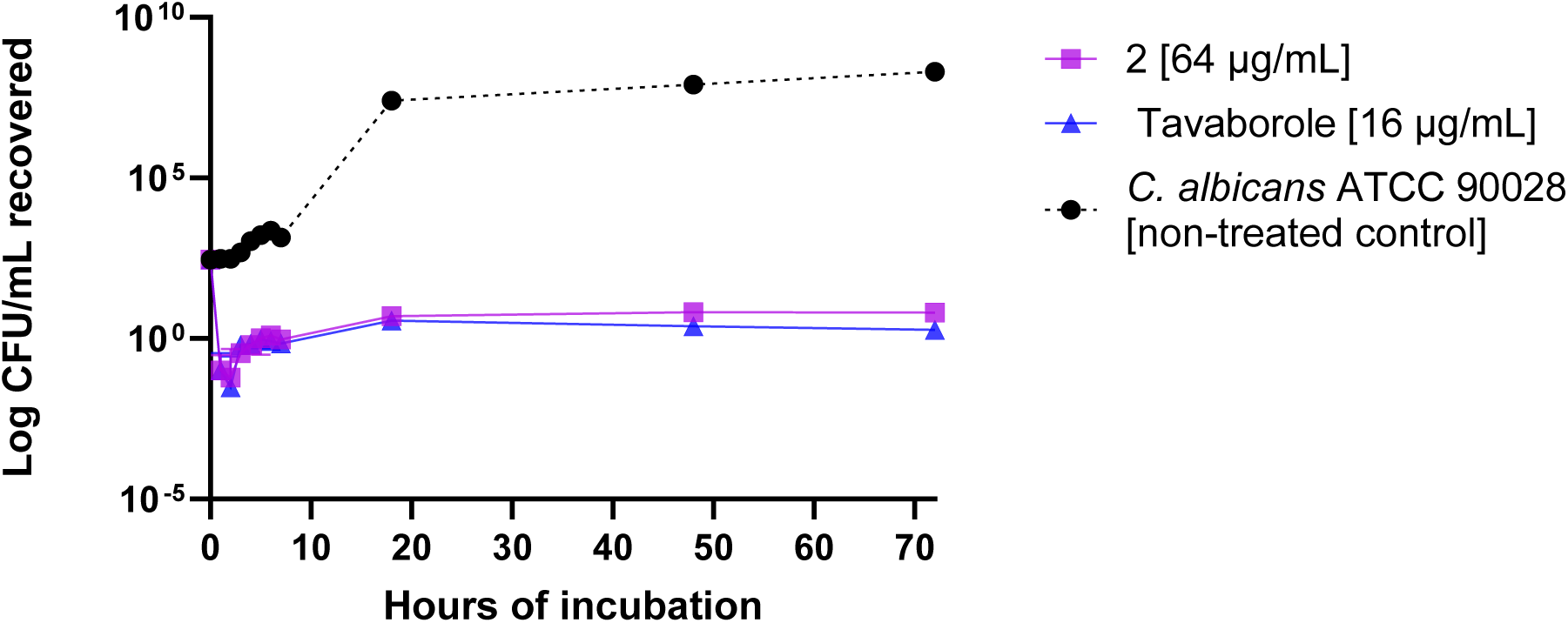
Time–kill curves of benzoxaborole derivatives against *Candida albicans*. *Candida albicans* 90028 time-kill kinetics assays. Comp. **2** (64 µg/mL) and **Tavaborole** (16 µg/mL) were added to the Sabouraud medium containing the blastoconidia starting culture. 90028 growth is included as a control. Tavaborole was used as a reference. The log CFU/mL for all groups is determined at time 0 and at subsequent time points up to 70 hours. Statistically significant differences were observed between growth control and **2** or **Tavaborole** (P<0.05).

In **Fig. 3**, BIC_50_=2 μg/mL (biofilm-inhibiting concentration) is the lowest concentration of ref. that presented 50% inhibition of biofilm metabolic activity, when compared to the non-treated control. Treatments with 32 μg/ml of **2** inhibited biofilm metabolic activity by approximately 50% after 48 h (**Fig. 3**). Overall, as described below the results for microscopy, **2** displayed poor ability to inhibit morphogenesis in *C. albicans*.

**Fig. 3.**
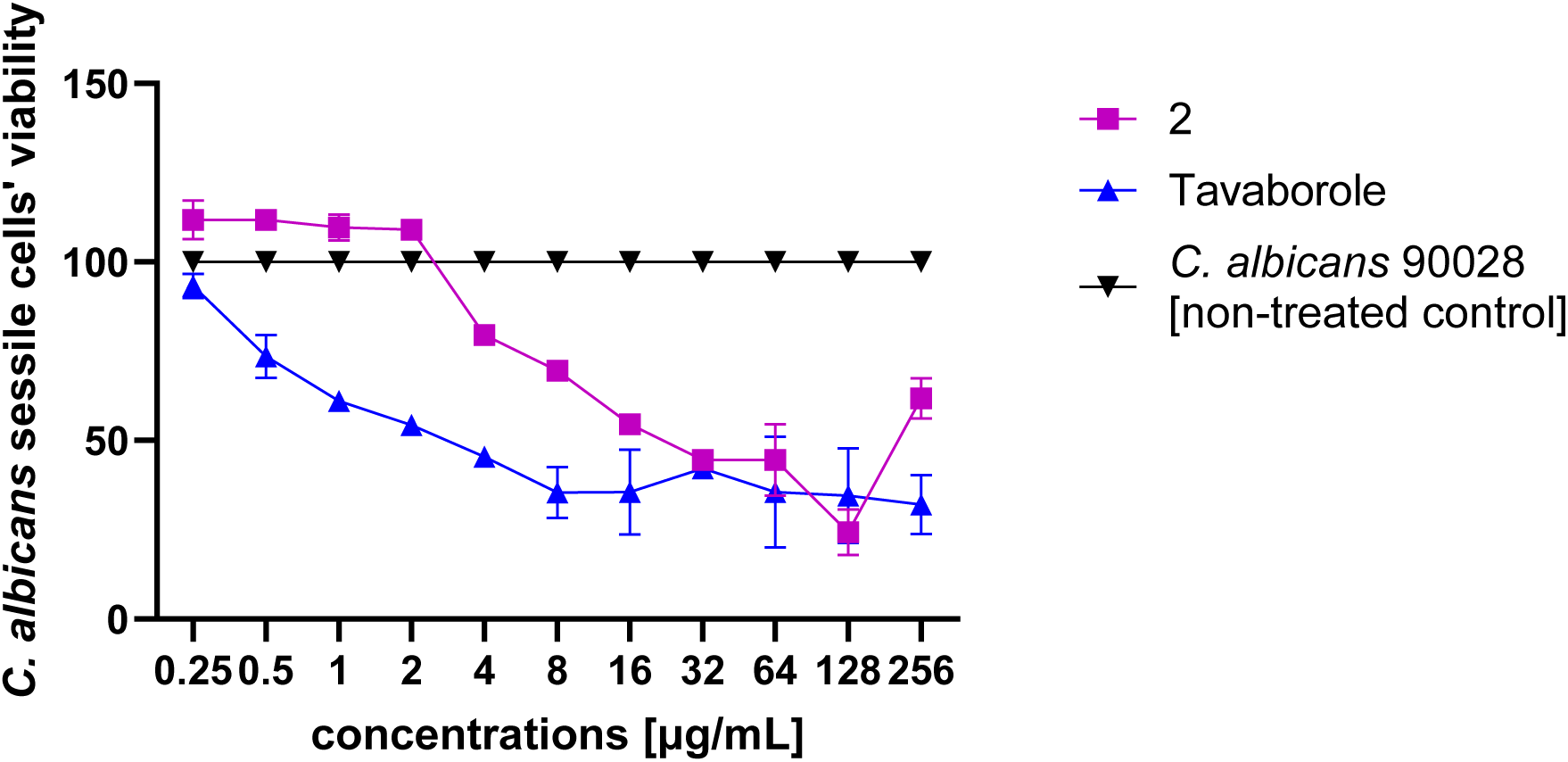
Biofilm inhibition by benzoxaborole derivatives The *Candida albicans* 90028 biofilms growing in 96-well plates and treated with **2** or **Tavaborole** for 18 h at 37°C and then analyzed by the MTS assay. The reduction of sessile cell viability was calculated by the following formula: %alive = OD490 test sample/OD490 control x 100%, control means cells treated with 0.96% DMSO. Data were presented as mean percentage ± SD for three independent repetitions. Multiple unpaired t-tests: non-treated control vs **2** (at 0.25 μg/ml, P<0.02); vs **Tavaborole** (at 0.25 μg/ml, P<0.03).

### Benzoxaboroles’ cytotoxicity *in vitro* and *ex vivo*

We evaluated cell viability in VERO E6 and PBMC with a 3-(4,5-dimethylthiazol-2-yl)-5-(3-carboxymethoxyphenyl)-2-(4-sulfophenyl)-2H-tetrazolium (MTS) reduction assay. The cytotoxic effect of **2** and **Tavaborole** (ref.) was assessed. The results are presented in **Fig. 4** and the degrees of viability were determined as the IC_50_ value of the viable cells. IC_50_ values were > 128 μg/mL for **2**. Ref. showed a weak cytotoxicity to VERO E6, at 256 μg/mL approximately 80% of cells were viable indicating a higher selectivity of the ref-drug to the pathogen than to the mammalian cells. Contrariwise, ref. and **2** showed IC_50_ at 2 μg/mL against PBMCs. Both **2** and ref. displayed selectivity in cytotoxic effects.

**Fig. 4.**
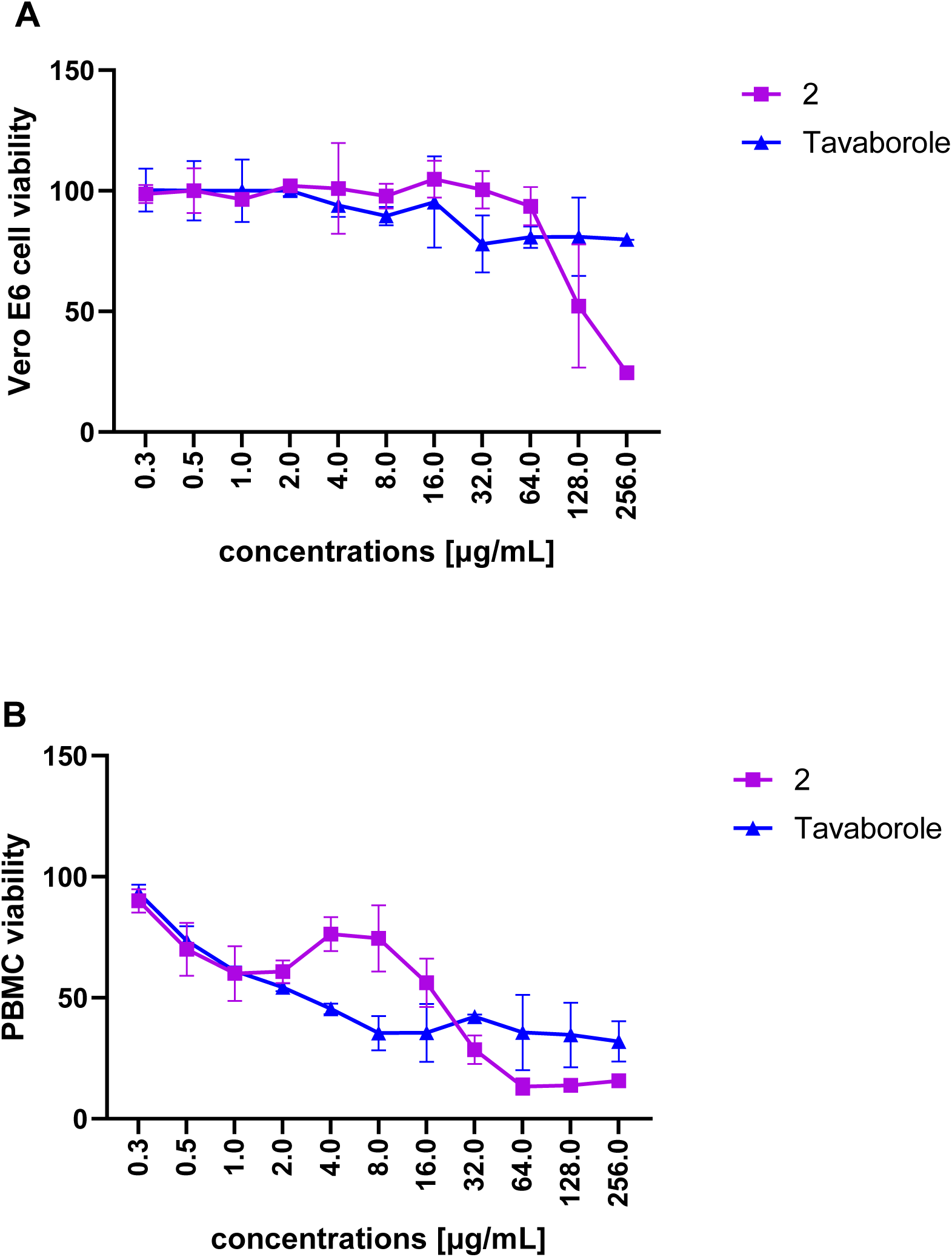
Cytotoxicity studies of benzoxaboroles against eukaryotic Vero E6 and PBMC cells Eukaryotic cells: Vero E6 (A) and PBMCs (B) were grown in 96-well plates and treated with **2** or **Tavaborole** for 18 h at 37°C and then analyzed by the MTS assay. The cell viability was calculated by the following formula: %alive = OD490 test sample/OD490 control x 100%, control means cells treated with 0.96% DMSO. Data were presented as mean percentage ± SD for three independent repetitions. (A) unpaired t-test: Tavaborole vs **1** was significantly different (P=0.0196) and Tavaborole vs **2** was significantly different (P=0.0035). (B) unpaired t-test: Tavaborole vs **2** was statistically insignificant (P=0.2763).

### Embryotoxicity of benzoxaborole derivatives

*In vivo,* the toxicity of benzoxaborole derivatives was investigated according to OECD Guidelines for The Testing of Chemicals (Fish Embryo Acute Toxicity (FET) Test). Compound **2** exhibited significantly lower embryotoxicity compared to **Tavaborole** (ref.). Both studied agents caused no or minimal mortality. Tavaborole at the highest concentration (4 µg/mL) caused significant (P=0.0229) zebrafish larval mortality during 96 h of exposure (**Fig. 5A**). However, Tavaborole significantly decreased the hatch rate. At the highest concentration studied (4 µg/mL), it completely prevented hatching (P<0.001), while at 2 µg/mL, it reduced hatching to 53.3±6.33% (P<0.0001) at 96 hpf (hours post fertilization). In contrast, **2** had no significant effect on hatching, and in each studied concentration 100% of the embryos hatched (**Fig. 5B**). Morphological analysis revealed that **2** at lower tested concentrations had minimal impact on the zebrafish embryos and larvae, except for the delay in swim bladder inflation. However, at the highest concentration tested (4 µg/mL), it caused notochord bending and mild pericardial edema. Minor notochord bending was sporadically observed for **2** at 2 µg/mL. In contrast, ref. induced morphological changes even at the lowest tested concentration, with the severity of changes correlating with concentration. The effect was visible starting from 24 hpf (data not shown) but was best expressed at 96 hpf. Pericardial edema was observed at all tested concentrations of ref., while higher concentrations (1-4 µg/mL) also led to tail and head deformities, small eyes, notochord bending, and yolk sac edema, lack of swim bladder inflation (**Fig. 6**). These findings suggest that **2** is much less toxic to zebrafish embryos compared to ref., since ref. even at the lowest concentrations tested disrupts the development of zebrafish.

**Figure 5.**
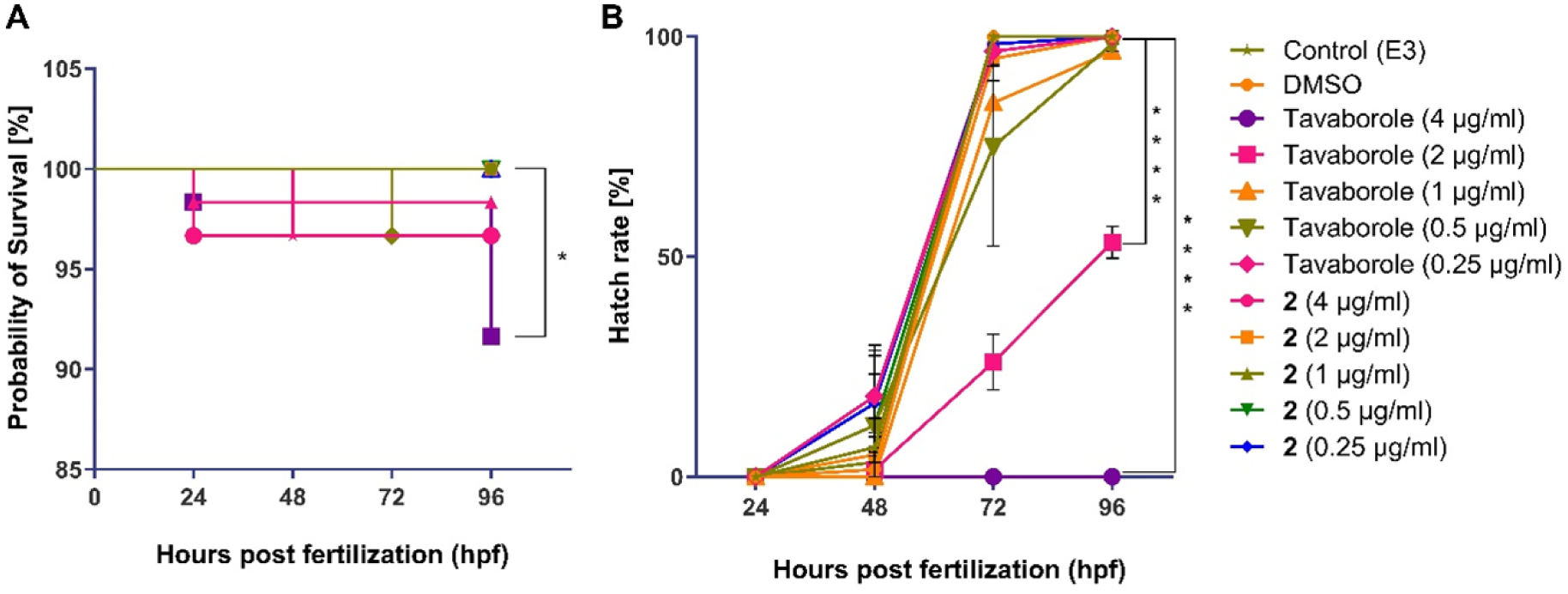
Probability of survival and hatch rate of zebrafish embryos after exposition to **Tavaborole** and **2** (A) Kaplan-Meier Survival Curve for Tavaborole and compound **2**. n=60, obtained from 3 independent repetition, *p<0.05 – Control vs Tavaborole (4 µg/ml; Log-rank (Mantel-Cox) test). (B). Hatching rate for Tavaborole and compound **2**. Data presented as mean ± SEM, n=3. For each repetition, the percentage was calculated from 20 individuals. p<0.0001 – Control vs Tavaborole (2 and 4 µg/mL; One-way ANOVA followed by Dunnett’s multiple comparisons test.

**Figure 6.**
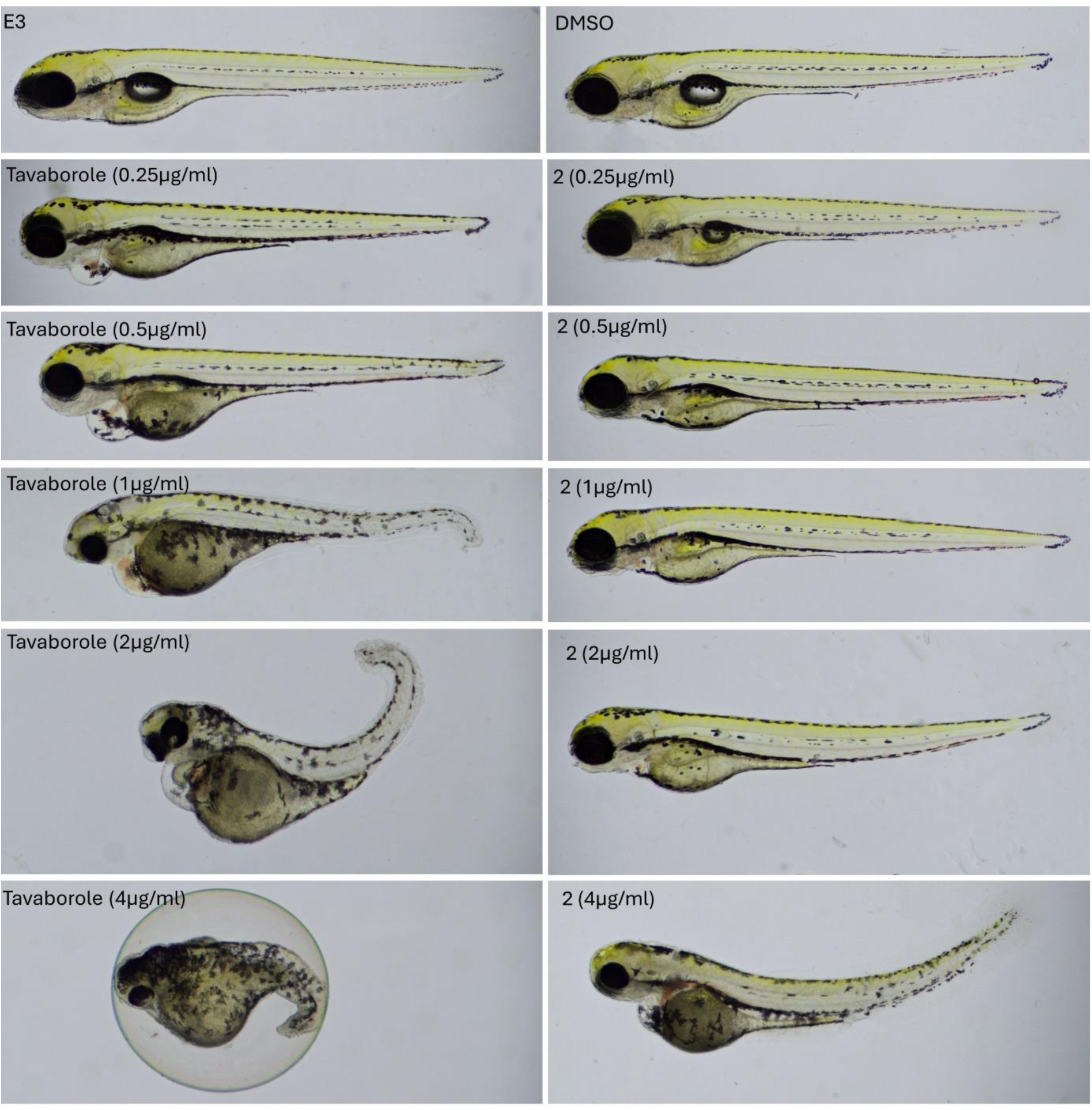
Zebrafish larvae 96 hours post fertilization exposed to **Tavaborole** and **2** E3 – not treated reference, DMSO – in DMSO, Tavaborole caused morphological changes even at the lowest tested concentration. Pericardial edema is evident across all Tavaborole concentrations, while higher concentrations also result in other malformations. Compound **2** has minimal impact on the development of zebrafish embryos and larvae at lower concentrations. At the highest concentration, it induces notochord bending and mild pericardial edema.

### Insights into cell membrane and cell wall composition

Encouraged by the above-described results obtained for **2**, insights into the cell membrane and cell wall composition were made. After 18 h of incubation with **2**, the *C. albicans* biofilm structure was not affected (**Fig. 7** WT+C). In contrast, *cnb1Δ/cnb1Δ* manifested a lack of filaments without the benzoxaborole treatments (**Fig. 7** Δ/Δ). In the test *C. albicans* biofilm treated with **2** (**Fig. 7** WT+C), the presence of necrotic fibroblasts (PI red fluorescence) was comparable with untreated biofilms (**Fig. 7** WT).

**Figure 7.**
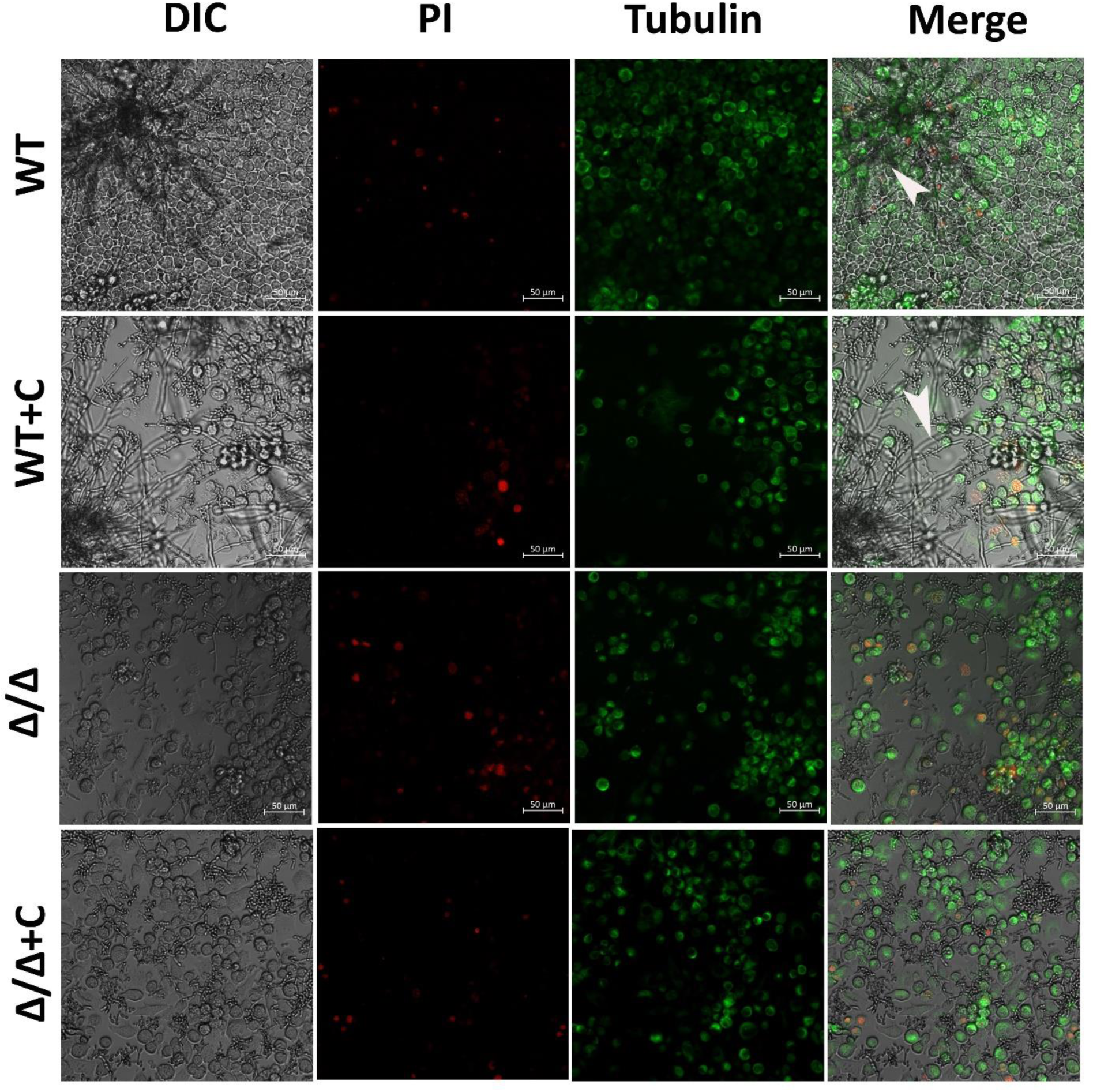
Confocal laser scanning microscopy images of the *Candida albicans* biofilms grown on fibroblasts after 18 h in the presence (test) and absence (control) of **2**. Live imaging of co-culture of L929 fibroblasts (labeled green with Tubulin tracker) and *Candida albicans* sessile cells. Labeling of the monolayer of fibroblasts with propidium iodide (PI) fluorophore allows the monitoring of fungal invasion (hyphal growth) of the wild-type *C. albicans* and lack of hyphae of *cnb1Δ/cnb1Δ.* The probes label the nucleus and cytoskeletal proteins (tubulins) in real-time. The antifungal agent does not inhibit *C. albicans* morphogenesis during invasion L929. DIC means differential interference contrast, PI - propidium iodide, tubulin means cytoskeletal protein stained with Tubulin tracker, Merge means DIC and fluorescence merged, WT means wild-type of C. albicans, Δ/Δ means *cnb1Δ/cnb1Δ*, WT+C means wild-type treated with **2** at 16 µg/mL, Δ/Δ + C means *cnb1Δ/cnb1Δ* treated with compound **2** at 16 µg/mL. Bar scale 10 μm.

### Differential expression analysis of control versus treated *Candida albicans.* Gene Ontology (GO) terms enrichment analysis of DEGs

The RNA-Seq of *C. albicans* control and bis(benzoxaborole) **2** treated replicates produced 31,265,125 reads on average. Each replicate’s reads are uniquely mapped to the reference genome with > 93 % on average (**Table S1**). The RNA-Seq read alignments were used to calculate gene expression count. Then, the gene expression count matrix was used to calculate differential expression to compare gene expression among the control vs treated group. The genes differentially expressed are in blue (**Figure 8**), and these differentially expressed genes (DEGs) are filtered with P ≤ 0.05 to be considered significant DEGs and studied for further downstream analysis. The significant DEGs are also summarized as up and down-regulated based on their log fold change (**Table S2**), and the top 30 significantly differentially expressed genes (by padj values ≤ 0.05) are shown in the plot (**Figure S2**).

**Figure 8.**
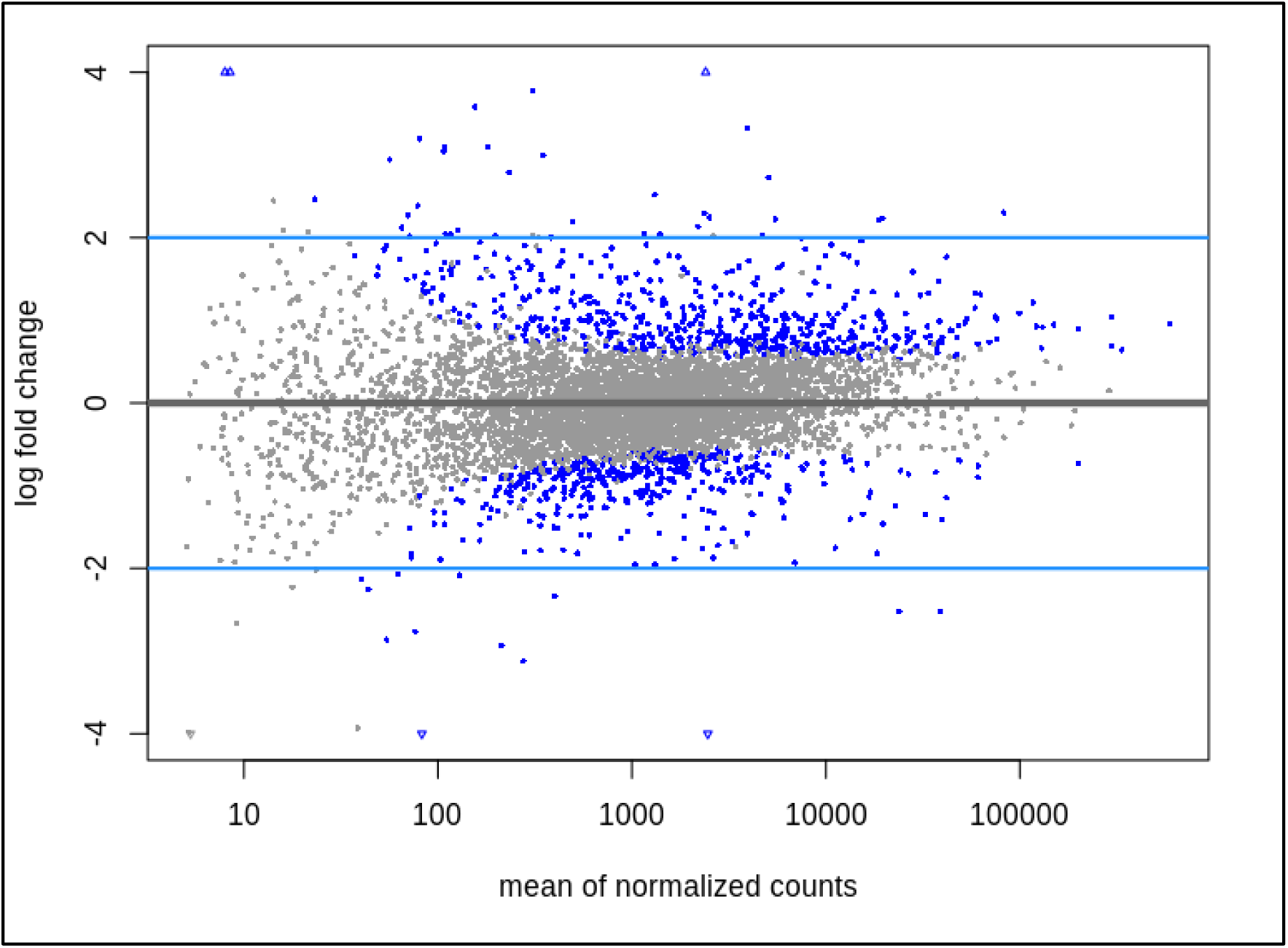
Global view of the differentially expressed genes

A total of 441 genes were upregulated, indicated by a positive log2 fold change, suggesting an increased expression level in the treated condition compared to the untreated condition. Conversely, 279 genes were identified as downregulated, denoted by a negative log2 foldchange, indicating a decreased expression level in the treated condition compared to the untreated condition. The global view of top differentially regulated genes is shown using a volcano plot (**Figure 9**), where the log-transformed adjusted p-values are plotted on the y-axis and log2 fold change values on the x-axis.

**Figure 9.**
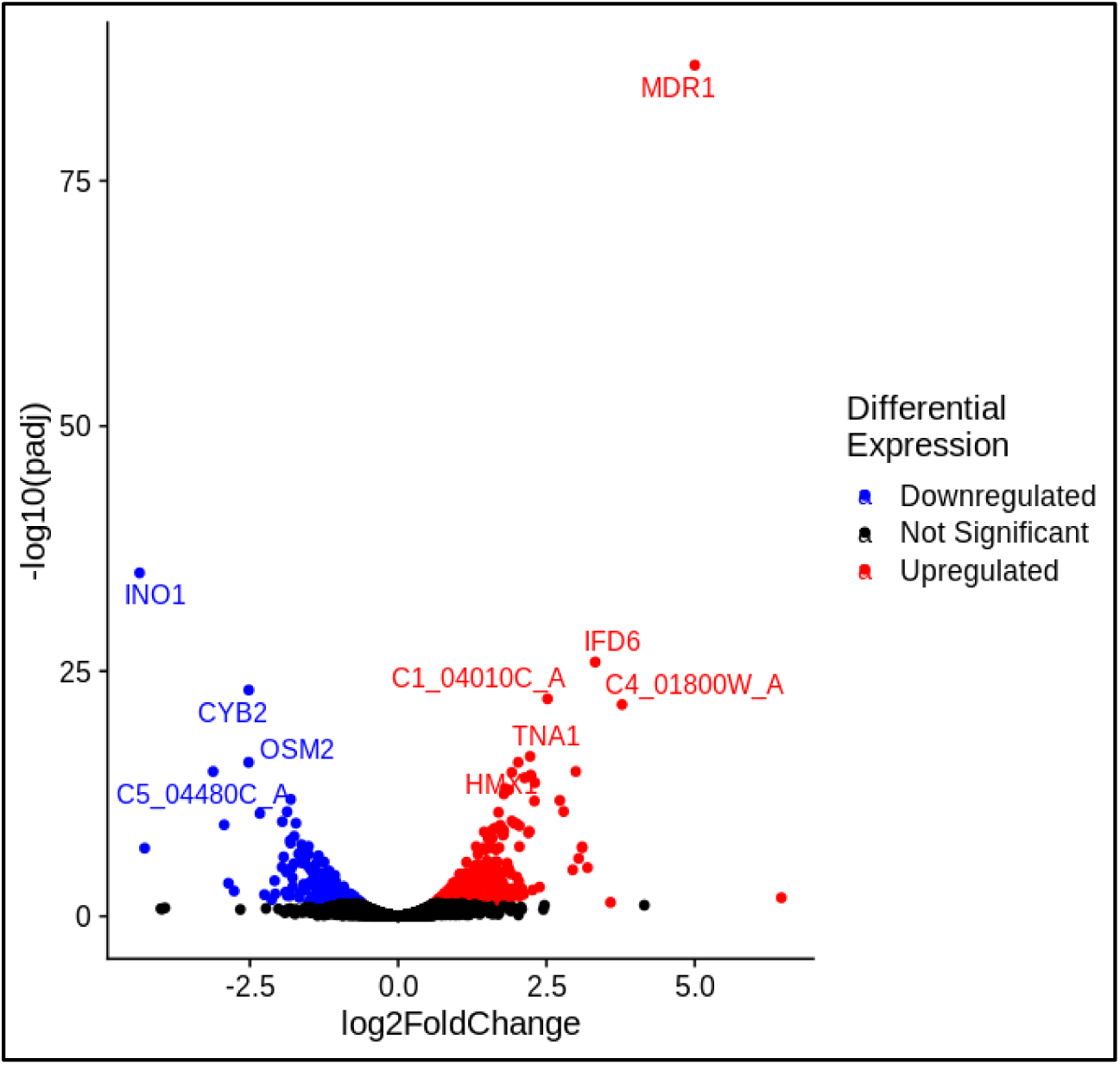
The volcano plot maps fold changes against p-adj and highlights the set of significantly differentially expressed genes. Upregulated significant genes (log2FoldChange > 0 and padj < 0.05) are red, downregulated significant genes (log2FoldChange < 0 and padj < 0.05) are blue and Not Significant are > 0.05.

**Figure 10.**
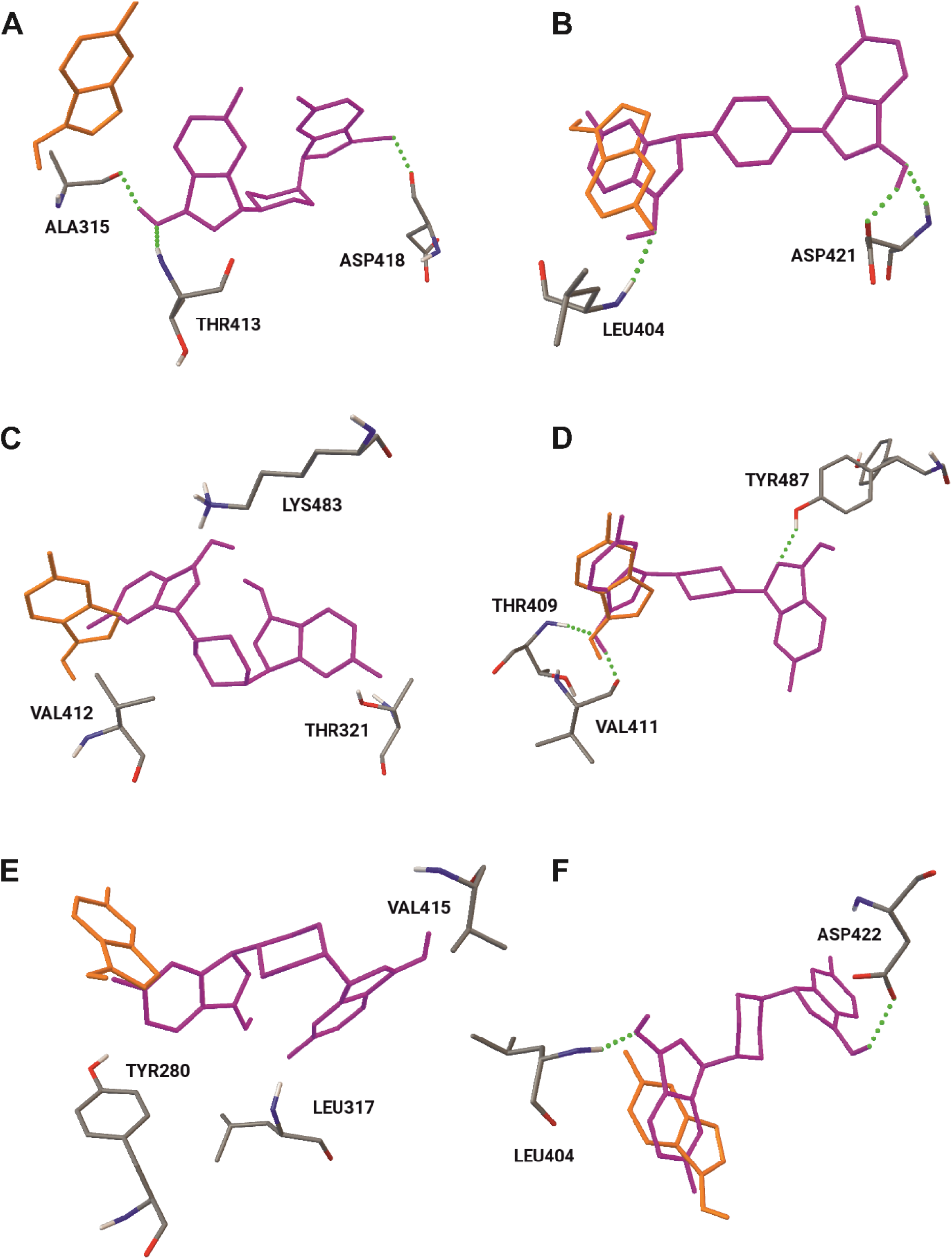
Structures of Tavaborole and stereoisomers of **2** docked into the active site of *C. albicans’* LeuRS Structures of Tavaborole (orange) as well as stereoisomers of **2** (pink, A: ***R,R*-2**, B: ***R,R*-2-a,** C: ***S,S*-2,** D: ***S,S*-2-**a, E: ***R,S*-2**, F: ***R,S*-2-a**) docked into the active site of *C. albicans’* LeuRS. Green dotted lines stand for hydrogen bonds. Structure ***S,S*-2-**a (D) shows the closest arrangement to the reference drug (Tavaborole).

The genes produced in this DEG analysis have exhibited diverse expressions upon treatment conditions. The Gene Ontology [29] association analysis was conducted to find the association of Gene ontology terms such as biological process (BP), molecular function (MF), and cellular components (CC) and corresponding functional descriptions to the respective DEGs. The top 10 most significant DEGs are associated with cellular components and molecular function GO terms, and most are upregulated (**Table S3**); a complete list is in Supplementary **Table S1**. The GO enrichment analysis [30] reported several observations: a) All the terms were enriched by g:Profiler in the GO resources[29], where 25 GO terms were enriched in Molecular function, 127 in biological processes, and 14 in the cellular component, respectively (**Figure S4**, **Table S2**). As mentioned in Methods, two more stages were employed to pick highlighted GO terms, which can be seen with numeric IDs in **Figure S4** and their details in **Table S4-6**.

The postulated molecular mechanism of Tavaborole’s anti-*C. albicans* action is based on the crystal structure of the LeuRS reported in 2015.[31] The experimental structure has been successfully reproduced in docking studies, enabling the determination of the inhibition constant of the Tavaborole-AMP adduct for comparative purposes.[32] Further docking studies showed that Tavaborole’s analogues including bis(benzoxaboroles) may follow a similar mechanism of action.[21],[33] The Current study covers not only docking studies of the AMP adducts of the boronic species but also the native compounds which is Tavaborole and compound **2**. The whole protein molecule has been included in the docking procedure and the position of the AMP-Tavaborole adduct in the crystal structure taken from literature was considered as the active center of the enzyme as well as the starting point of all the docking procedures. **Table 2** shows the determined inhibition constants of the lowest energy structures docked into the active site of the enzyme as well as the list of amino acids producing hydrogen bonds with studied boronic molecules. It is worth noting, that all the determined inhibition constants of **2** are lower than those of Tavaborole, nevertheless, they all are within the micromolar range, denoting rather weak interactions (**Table 2**). The docking procedure identified also several structures of slightly lower inhibition constants, however all outside the active site (**Table S7**). Most of the docked-in structures within the active site of LeuRS, adopt different positions in comparison to the Tavaborole molecule, which is however not the case for isomer ***S,S*-2-a** which forms hydrogen bonds with Thr409 and Val411, the same as Tavaborole does.

**Table 2.** Inhibition constants as well as hydrogen-bonded amino acids of the lowest energy structure of Tavaborole and stereoisomers of **2** docked into the active site of *C. albicans’* LeuRS.

|  | Inhibition constant | Amino acids forming hydrogen bonds<br>with the ligand |
| --- | --- | --- |
| **Tavaborole** | 989.25μM | THR409, VAL411, GLY410, |
| *R,R*-**2** | 5.13μM | ALA315, VAL412, ASP418 |
| *R,R*-**2-a** | 12.89μM | LEU404, ASP421, ASP421 |
| *S,S*-**2** | 2.24μM | no hydrogen bonds |
| *S,S*-**2-a** | 6.45μM | THR409, VAL411, TYR487 |
| *R,S*-**2** | 9.13μM | no hydrogen bonds |
| *R,S*-**2-a** | 32.55μM | LEU404, ASP422 |

Docking studies covered also the adducts of Tavaborole as well as all the stereoisomers of **2** with adenosine monophosphate (AMP), strictly according to the previously reported procedure.[32] Table 3 shows exclusively results for the structures docked into the active site of the *C. albicans* LeuRS enzyme. Most of the determined inhibition constants are much lower in comparison with those of native ligands presented in Table 2 and are in the nanomolar range (Table 3). Nevertheless, none of the **2-AMP** adducts adopts a position analogous to that of Tavaborole-AMP (Figure 11, orange vs pink structures). None but one structure, namely ***R,S*-2-a-R-AMP-1** displays an inhibition constant lower than that of Tavaborole-AMP (Table 3). It is worth noting that both the configuration of **2** as well as arrangement of the AMP part of the adduct, strongly influence the binding in terms of both the binding part of the enzyme as well as the value of the inhibition constant.

**Figure 11.**
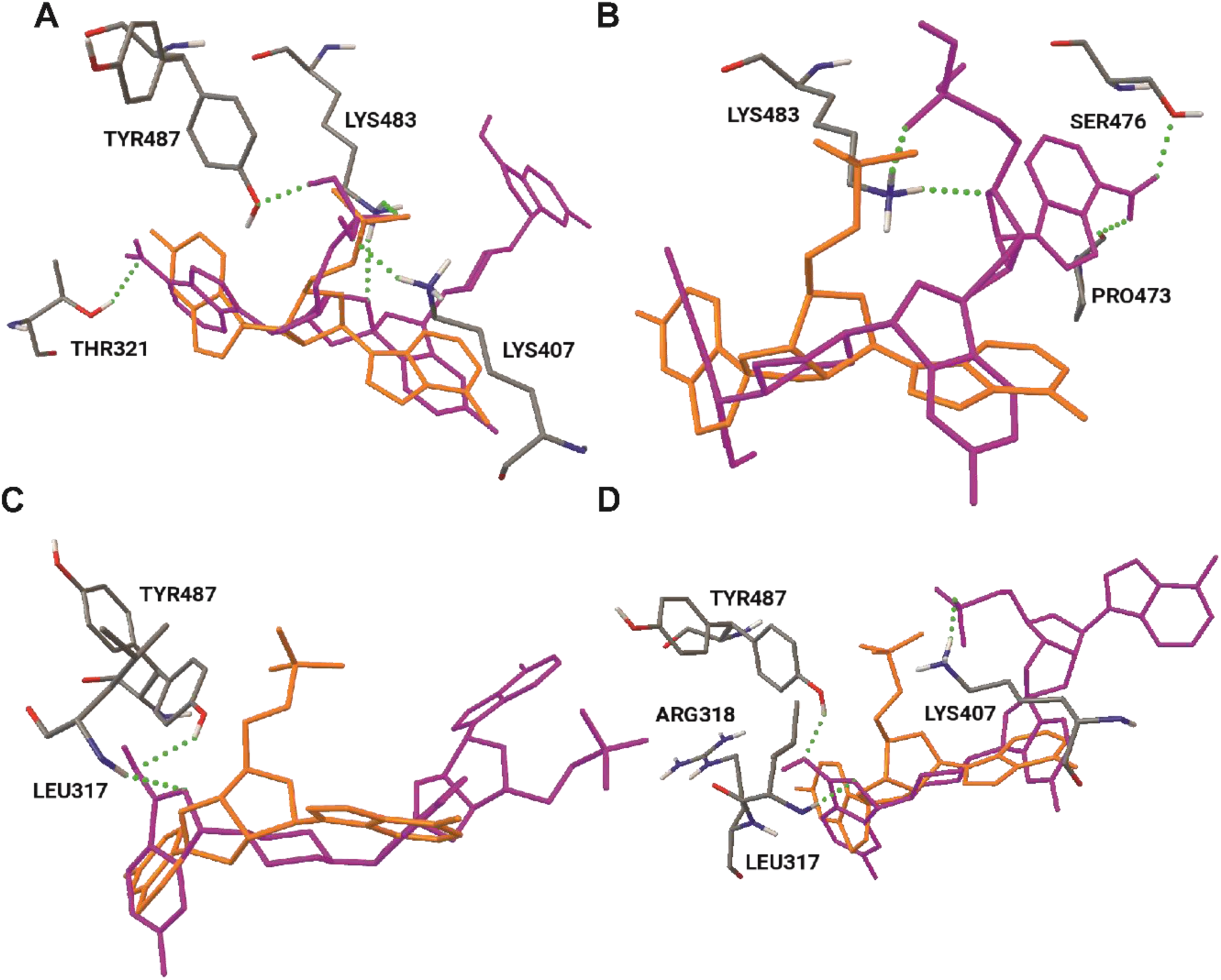
Structure of Tavaborole-AMP (orange) as well as of selected stereoisomers of **2-AMP** (pink) docked into the active site of *C. albicans’* LeuRS. Structures of Tavaborole-AMP (orange) as well as stereoisomers of **2-AMP** (pink, A: ***R,R*-2-a-AMP-1,** B: ***R,R*-2-a-AMP-2,** C: ***S,S*-2-a-AMP-1,** D: ***S,S*-2-a-AMP-2**) docked into the active site of *C. albicans’* LeuRS.

**Figure 12.**
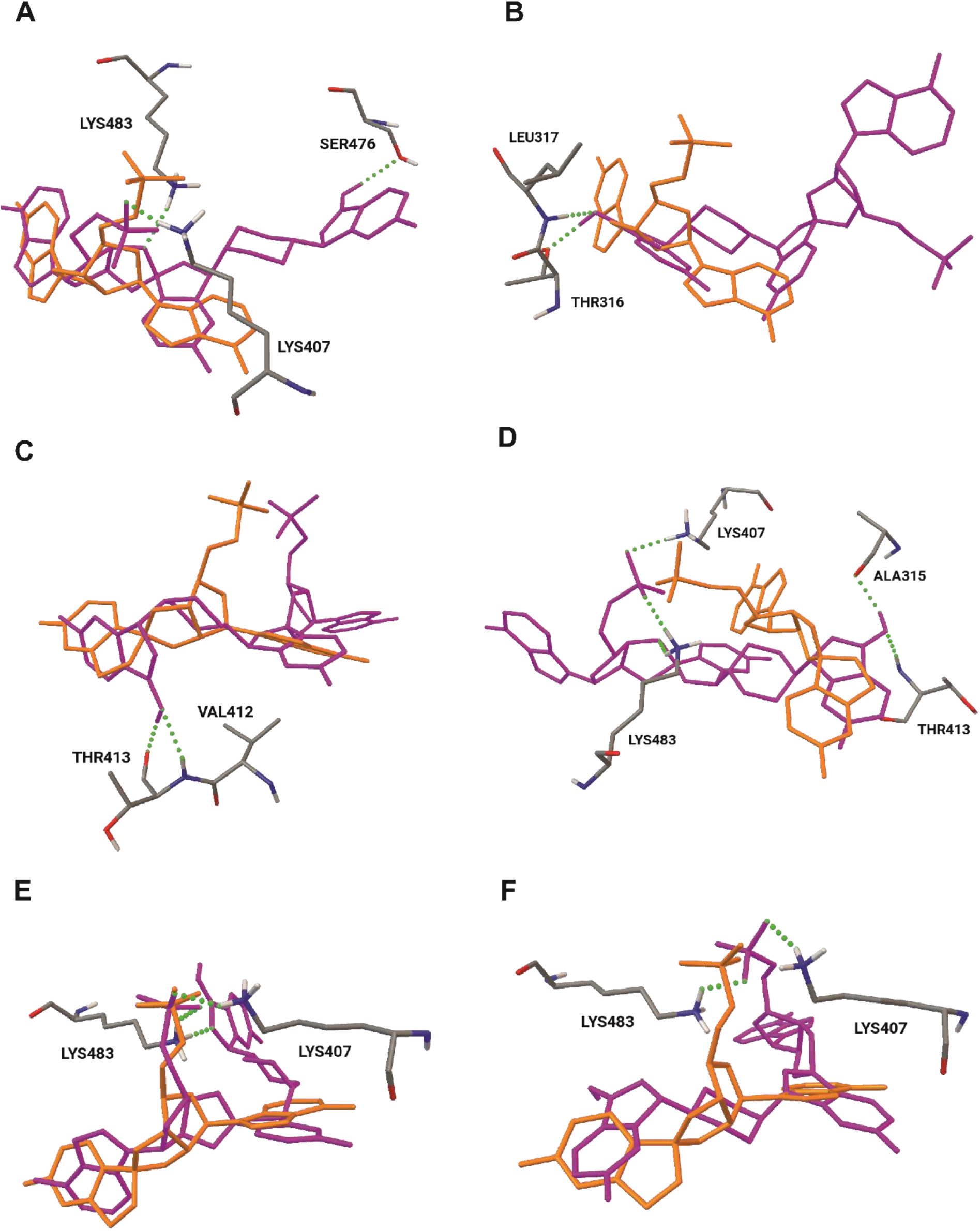
Selected structures of Tavaborole-AMP as well as of stereoisomers of **2-AMP** docked into the active site of *C. albicans’* LeuRS. Structures of Tavaborole-AMP (orange) as well as stereoisomers of **2-AMP** (pink, A: ***R,S*-2-*R*-AMP-1**, B: ***R,S*-2-*S*-AMP-1**, C: ***R,S*-2-*S*-AMP-2**, D: ***R,S*-2-a-*S*-AMP-2**, E: ***R,S*-2-a-*R*-AMP-1**, F: ***R,S*-2-a-*R*-AMP-2**): docked into the active site of *C. albicans’* LeuRS.

**Table 3.** Inhibition constants as well as hydrogen-bonded amino acids of the lowest energy structures of the adenosine monophosphate (AMP) adducts with Tavaborole and stereoisomers of **2** docked into the active site of *C. albicans’* LeuRS.

|  | Inhibition<br>constant | Amino acids forming hydrogen bonds<br>with the ligand |
| --- | --- | --- |
| Tavaborole-AMP[32] | 1.93nM | LEUS317, LEU404, LYS407, LYS483, LYS483 |
| **R,R-2-AMP-1** | 377.22nM | LYS407, LYS483, SER476 |
| **R,R-2-a-AMP-1** | 4.38nM | THR321, LYS407, LYS483, TYR487 |
| **R,R-2-a-AMP-2** | 671.5nM | PRO473, SER476, LYS483 |
| **S,S-2-a-AMP-1** | 17.41μM | LEU317, TYR487 |
| **S,S-2-a-AMP-2** | 101.79nM | LEU317, LYS407, TYR487 |
| **R,S-2-R-AMP-1** | 17.27nM | LYS407, SER476, LYS483 |
| **R,S-2-S-AMP-1** | 7.99μM | THR316, LEU317 |
| **R,S-2-S-AMP-2** | 53.46nM | VAL412, THR413 |
| **R,S-2-a-R-AMP-1** | 1.68nM | LYS407, LYS483, LYS483 |
| **R,S-2-a-R-AMP-2** | 9.45nM | LYS407, LYS483 |
| **R,S-2-a-S-AMP-2** | 80.22nM | ALA315, LYS407, THR413, LYS483, LYS483 |

## Discussion

The benzoxaborole concentration-response relationship described the important lack of cytotoxicity/ embryotoxicity between agents’ concentrations and anti-*C. albicans* effect. This study provided new information on the pharmacodynamic relationship between benzoxaborole concentrations and blastoconidial population dynamics. The individual analysis of compounds indicated that **2** and **Tavaborole** (ref.) showed comparable effects on *C. albicans*. Benzoxaboroles inhibited the growth during 72 h, the killing activity started after 2 h of that time (**Fig. 2**). We observed metabolic changes after exposure to benzoxaborole (%R=64) in **Fig. 1A**. Time-kill kinetics assay helps to understand interactions that exist between *C. albicans* and benzoxaborole. Benzoxaborole derivatives were established through the time-kill kinetics study to be fungistatic. In conclusion, the time–kill kinetic assays revealed that **2**’s activity against *C. albicans* was comparable to ref. However, we demonstrated that the activity of both compounds tested was moderate thus the further concentration effectiveness *in vivo* needs to be assessed in the future. Based on our findings using the zebrafish model (**Fig. 5 and 6**), we indicate that tested compounds’ optimization challenging to satisfy all requirements (safety and fungicidal activity) is still in high demand. The *C. albicans* pathogenic mechanism presented in biofilm growing on fibroblasts was studied *in vitro* using co-culturing in a static biofilm model (**Fig. 7**). This biofilm model was used to study the interactions between the *C. albicans* morphogenesis and benzoxaboroles. Biofilm formation reduces benzoxaborole penetrating abilities (**Fig. 7**). *Candida albicans* living in biofilm (**Fig. 3** and **Fig. 7** WT+C) developed resistance (development of invasive hyphae) to benzoxaborole derivatives compared with those existing in free-living forms (planktonic forms in **Fig. 1A** and **Table 1**). Subsequently, working with Confocal Laser Scanning Microscope (CLSM) suggested the existence of hyphae under the benzoxaborole treatment. On the contrary, our results point out the benzoxaboroles’ ability to repress the metabolism of sessile cells (**Fig. 3**) by the alternation of MDR efflux pumps and their coactivators (**Fig. 8** and **9**, **Table S3**). We found decreased expression of the fumarate reductase gene (*OSM2* in **Fig. 8** and **Table S3**) involved in the yeast-to-hyphae transition [34]. Moreover, the importance of ergosterol in the *C. albicans* resistance to benzoxaborole was tested. Our results (**Fig. 1B**) clearly illustrate that benzoxaborole antifungal development targets ergosterol binding. In line with that membrane mechanism, RNAseq data indicated that efflux pumps in *C. albicans* were upregulated (**Fig. 8** and **9**, **Table S3**). The cellular detoxification based on the efflux pumps encoded by the *MDR1* gene [35] was employed in exporting benzoxaborole to the exterior of the *C. albicans* cells. Contrariwise, *INO1* (Inositol-1-phosphate synthase) was repressed under the benzoxaborole treatment (**Table S3**). This gene is responsible for inositol synthesis, it is the growth factor supporting the formation of glycophosphatidylinositol (GPI)-anchored glycolipids of *Candida* and hence pathogenesis [36]. We found that *MDR1* expression by benzoxaborole **2** was accompanied by the *IFD6* (aldo-keto reductase) increase. The studies on benzoxazole derivatives revealed a complex picture where the induction of drug efflux pump expression (*MDR1*) requires the coordination of multiple coactivators (*IFD6, TNA1* encoding putative nicotinic acid transporter). We showed that benzoxaboroles represent a similar resistance mechanism to azoles due to the subsequent expression of *MDR1* and *IDF6* [37]. This suggests that the effect of benzoxaboroles on efflux pump activity can be associated with changes in the ergosterol synthesis pathway [38]. Another interesting finding was that *C1_04010C_A* (encoding protein with a NADP-dependent oxidoreductase domain) appears to increase the overall fitness of *C. albicans* when challenged with benzoxaboroles (**Fig. 8**, **Table S3**). In line with our findings, recent studies by Bandara *et al*. [38] have also shown an increase in *C. albicans*’ antifungal resistance via upregulated proteins associated with drug resistance and virulence.

The noted variations in factors involved in various metabolic pathways, transport mechanisms, response to stress, formation of the external cell layer, enzymatic processes, and molecule binding processes indicate searching pathways for benzoxaborole sensitivity-restoring mechanism. Since the inhibition of this efflux activity may be performed in different ways, we showed that benzoxaboroles alternate the expression of efflux pumps (*MDR1*) and significantly downregulate well-characterized genes linked to virulence, such as *INO1, CYB2,* and *OSM2*. The enriched molecular function performed by DEGs (**Fig. S4**) indicates the pivotal role of oxidoreductase activity in response against **2**. Additionally, small molecule binding and transporter (aminoacylotRNA) activity are noteworthy to be mentioned as a target for benzoxaboroles. While new inhibitors’ design is often turned to the accessory sites such as the amino acid, ATP, or tRNA binding pockets [39], our docking studies confirmed the proposed interactions of benzoxaborole adenosinemonophosphate adduct with LeuRS. Moreover, *C5_04480C_A* was negatively regulated in response to the benzoxaborole stress. This gene is responsible for cell wall biogenesis, protein folding, modification, and destination[40]. Thus, our findings contribute to further understanding of molecular mechanisms regulated by benzoxaborole action mode. Concerning cellular components, cell surface, and external encapsulating structure are highly significant, indicating the involvement of genes in extracellular interactions with the stress generated by benzoxaborole. The core microcolony gene (*TNF1*) was upregulated significantly under the benzoxaborole treatment. The latter closely reflects a lack of total inhibitory activity of these agents against pathogenic forms of biofilm [41].

Given all the above, a potential mechanism for benzoxaboroles is the efflux pump inhibitors (EPIs) targeted into the reduced metabolism of sessile cells. Such an action has been previously reported for other phenylboronic species, yet not for benzoxaboroles [42],[43]. These conclusions are largely based on our transcriptomic data and experiments; therefore, appropriate functional assessments are necessary to verify these claims. Further investigations on sterol analyses and changes in plasma membrane composition are necessary to confirm this hypothesis. Since efflux pumps are considered important drug targets, benzoxaborole-altering efflux inhibitors can be important for the development of combination strategies in candidiasis. Our findings present innovative concept that can inspire further studies for designing and building new antifungal benzoxaborole.

## Experimental

### Docking studies

Molecular structures of stereoisomers of **2** were optimized in Gaussian03[44](b3lyp, [45] 6-311g(2d,p) [46],[47],[48],[49],[50],[51],[52],[53]. All the stereoisomers (*R,R*, *S,S* and *R,S*) were taken into account. The optimized structures were connected to the AMP molecule taken from crystal structure of the AN2690-AMP spiroboronate found in the crystal phase [31] to form the investigated ligands. Different structural patterns were assumed, varying in arrangement of the benzoxaborole phenyl and the adenosine rings in the resulting ligands. Structures denoted with “1” represent spiroboronates with those rings at the same side of the spiroboronate, and those denoted with “2” represent structures with rings at opposite part of the molecule. The resulting spiroboronates corresponding to all stereoisomers as well as the position of the adenosine rings were not optimized. The input structures of the AMP adducts have not been optimized. In case of every input structure a 300-step docking procedure according to the genetic algorithm has been carried out. The starting position of the ligand was analogous to that of Tavaborole known from the experimental complex[31]. The search for binding sites covered the whole protein structure.

### *Candida albicans* susceptibility testing

The percentage inhibition (%R) of each of the tested compounds was determined by broth microdilution in YEPD medium (A&A Biotechnology, Poland) at 30°C, according to the M27-A3 method[28] (CLSI guidelines). We performed manual broth microdilution with the concentration ranges from 0.5 to 64 µg/mL of YEPD. The stock solution of compounds (51,200 μg/mL) was prepared in DMSO (96%, Sigma-Aldrich). The starting cell suspension density was 1-3 x 10^2^ CFU/mL. The spectrophotometric measurements were taken after 18 and 48 h incubation at 625 nm by a microtiter plate reader Spark (Tecan Switzerland). The growth inhibition was calculated using the following formula: %R = 100 – (OD_625_ CTW – OD_625_ CTW)/(OD_625_ GCW – OD_625_ GCW). CTW means compound test wells, GCW growth control wells. The MFCs were determined as was described previously.[54] Appropriately, after 10-fold dilution series 100 μL aliquots from microtiter plates were used to inoculate YEPD agar plates (A&A Biotechnology, Poland) and after 48h incubation at 37°C the grown colonies were counted. Using the following formula: lgR = lgCFU/mL GCW – lgCFU/mL CTW, CFU/mL = (mean number of colonies grown on 3 plates) x (inverse of the dilution = 10^-4^) x 10 (counted per 1 mL), MFC being defined as the reduction viability record (R) was calculated. LgR ≥3 means fungicidal concentration, 0 ≤ lgR < 3 fungistatic, and 0 ≤lgR not active. Data were presented as mean percentage ± SD for three independent repetitions.

The metabolic activity inhibition of Tavaborole compounds against *C. albicans* ATCC 90028 planktonic cells was assessed using MTS test (3-(4.5-dimethylthiazol-2-yl)-5-(3-carboxymethoxyphenyl)-2-(4-sulfophenyl)-2H-tetrazolium, MTS, Promega, USA). The suspension of *C. albicans* cells in medium YEPD at density of 1-3 x 10^2^ CFU/mL were incubated with compounds **2** and Tavaborole at the conc. range of 0.25 to 512 μg/mL for 48 h at 37^°^C. Then 10 μL of MTS solution was added to each well and 2 h-incubation in the darkness was performed. The optical densities were measured at 490 nm using a microtiter plate reader Spark (Tecan, Switzerland). The proliferation was determined using the following formula: %alive = OD_490_ test sample/OD_490_ control x 100%, control was treated cells with 0.96% DMSO (comp. solvent). Data were presented as mean percentage ± SD for three independent repetitions.

The benzoxaborole derivatives’ interaction with exogenous ergosterol was determined. The suspension of *C. albicans* cells at density 1,74 x 102 CFU/mL was prepared in YEPD medium supplemented with ergosterol at final conc. of 400 μg/mL (prepared directly before the experiment in 10% DMSO v/v and 1% Triton X-100 v/v). The tested cells were treated with comp. in the range from 0.25 to 256 μg/mL, the control was treated with 0.96% DMSO and incubated for 48 h at 35oC. The viability measurements were taken at 405 nm by plate reader Spark Control M10 (Tecan Group Ltd, Germany) as described above. Data were presented as mean percentage ± SD for three independent repetitions.

### Time-Kill assay

For 3 days the activity of comp. **2** and Tavaborole (both at MFC conc.) was studied.[55] The experiment was performed using *C. albicans* suspension at 1-3 x 10^2^ CFU/mL in YEPD medium. As a control, the cells were treated with 0.96% DMSO. At the selected times, the cell suspension was inoculated onto the Petri dishes and incubated at 37°C for 48h, then the colonies were counted. The logarithm of the reduction viability record (logR) was determined as described above.

### Activity against preformed biofilm assay

Biofilm was formed on a 96-well microtiter plate using *C. albicans* ATCC 90028 suspension in YEPD medium at 1-3 x 10^2^ CFU/mL during 18 h incubation at 37^°^C.[56],[57] Preformed biofilm was treated with comp. **2** and Tavaborole in the range 0.25-512 μg/mL for 18 h at 37^°^C and then analyzed by MTS assay. The reduction of sessile cell viability was calculated by the following formula: %alive = OD_490_ test sample/OD_490_ control x 100%, control means cells treated with 0.96% DMSO (comp. solvent). Data were presented as mean percentage ± SD for three independent repetitions.

### Cytotoxicity *in vitro* and *ex vivo* assay

The toxicity effect of comp. **2** and Tavaborole in the range of conc. from 0.25 to 512 μg/mL was characterized using the kidney epithelial cell line Vero (ATCC CCL-81, LGC Poland) as previously described.[28] The VeroE6 cell monolayer was formed during the 18-h incubation period at 37°C and 5% CO_2_ using cell suspension at 1 x 10^6^ cells/mL MEM medium (Gibco) supplemented with 10% fetal bovine serum (Gibco), 1% antibiotics (Sigma-Aldrich). Subsequently, compounds were added to proper wells in triplicate and incubated for 18 h under the same conditions. 10 μL MTS solution was added after 2-h incubation in the darkness, the optical measurements were made at 490 nm. The percentage of alive cells was calculated as % alive = OD_490_ test sample/OD_490_ control x 100%, control was cells treated with 0.96% DMSO (comp. solvent). Data were presented as mean percentage ± SD for three independent repetitions. Unpaired t-test (P <.05 significantly different) was used to analyze data (mean values were analyzed).

PBMCs were isolated from human venous blood using Ficoll-Histopaque-1077 (Sigma) and gradient centrifugation.[58] PBMCs at density 1 x 10^6^ cells/mL medium Opti-MEM (Gibco) supplemented with 10% fetal bovine serum (Gibco), 1% antibiotics (Sigma-Aldrich), were incubated with **2** and **Tavaborole** during 18h at 37°C and 5% CO_2_ in 24-well plates. Subsequently, 10 μL MTS was added to each well and incubated for 2h in the darkness. The absorbance measurements were made at 490 nm using a microtiter plate reader Spark (TECAN Switzerland). The percentage of alive cells was calculated as described above. Data were presented as mean percentage ± SD for three independent repetitions. Unpaired t-test (P <.05 significantly different) was used to analyze data (mean values were analyzed).

### Embryotoxicity

The toxicity of comp. **2** and Tavaborole for selected conc. was tested according to the Fish Embryo Acute Toxicity (FET) Test protocol [59] using zebrafish embryos of AB line. The control contained 24 embryos and other groups (4 μg/ml 3,4 – dichloroaniline-positive control, 0.03% DMSO – solvent control, and compounds at selected concentrations) contained 20 embryos that were exposed to the solvent on a separate 24-well plate with an internal control group of n = 4, on each plate. Incubation mediums were changed daily. During 4 days of incubation, the mortality, hatching rate, and possible malformations were observed daily, on each embryo tested. The experiment was repeated 3 times.

### Insights into cell membrane and cell wall composition using confocal laser scanning microscope

The *C. albicans* cells (at density 1 x 10^7^ CFU/mL) were treated with **2** at a final conc. of 64 μg/mL and inoculated onto L929 fibroblasts (ATCC CCL-1) and at the same time stained with Tubulin tracker and propidium iodide (PI) fluorophore. Using *C. albicans* and calcineurin mutants allowed the monitoring of morphogenesis under benzoxaborole exposure.[60] The control was the fibroblasts treated with the *C. albicans* wild-type. After 18-h incubation at 37°C the samples were observed using a confocal laser scanning microscope (CLSM, Zeiss Axio Observer 7 with LSM 900, detector Airyscan 2). All experiments were performed in triplicate.

### RNA-Sequence analysis for differentially expressed genes

RNA-Seq reads were analyzed for a quality check using FastQC1 v0.12.0, and the reads with bases quality >=30 (Phread score) and read size >= 35 bp were kept using Cutadapt[61]. The reference genome of *C_albicans*_SC5314_A22_current_chromosomes.fasta was sourced from The *Candida* Genome Database.[62] The reference genome was converted into a haploid genome by extracting the chromosomes with haplotype A only, given that a diploid genome has two haplotypes, with each chromosome with suffixes A and B. Then, read sequences were aligned to this haploid genome using RNA-seq by Expectation Maximization (RSEM)4 with --paired-end --strandedness reverse option.

The rsem-calculate-expression command from the RSEM suite produced the gene expression count for all samples. Each sample’s matrix of all these counts was built using a custom R script for differential expression analysis. We identified differentially expressed genes (DEGs) using Deseq2[63] and filtered with an adjusted p-value ≤ 0.05 (Benjamini-Hochberg adjustment) to be considered statistically significant. The Deseq2 also calculated log fold change based on differentially expressed genes. The gene ontology and annotation data were acquired from the gene ontology resource6, gene ontology R library,[64] and detailed features for respective annotation from The *Candida* Genome Database [62] in November 2023. The g:Profiler[30] was employed for the enrichment profiling of DEGs. The data source version from Ensembl for the organism is GCA000182965v3, and for gene ontology, it is BioMart classes: releases/2024-01-17. The Benjamini-Hochberg FDR threshold of 0.05 is considered statistically significant. Further, among significant terms, g:Profiler provides highlight driver terms based on two stages: first, organizing important terms into sub-ontologies based on their relationships, and second, pinpointing key gene sets that drive other significant functions nearby.

## References

1. Zirkel, J.; Klinker, H.; Kuhn, A.; Abele-Horn, M.; Tappe, D.; Turnwald, D.; Einsele, H.; Heinz, W.J. Epidemiology of Candida blood stream infections in patients with hematological malignancies or solid tumors. Med. Mycol. 2012, 50, 50–55, doi:10.3109/13693786.2011.587211.

2. Ju, Y.; He, L.; Zhou, Y.; Yang, T.; Sun, K.; Song, R.; Yang, Y.; Li, C.; Sang, Z.; bao, rui; et al. Discovery of novel peptidomimetic boronate ClpP inhibitors with noncanonical enzyme mechanism as potent virulence blockers in vitro and in vivo. J. Med. Chem. 2020, 0, doi:10.1021/acs.jmedchem.9b01746.

3. Görkem, A.; Sav, H.; Kaan, Ö.; Eren, E. Coronavirus disease and candidemia infection: A case report. J. Mycol. Med. 2021, 31, 101155.

4. Benedict, K.; Jackson, B.R.; Chiller, T.; Beer, K.D. Estimation of Direct Healthcare Costs of Fungal Diseases in the United States. Clin. Infect. Dis. 2019, 68, 1791–1797, doi:10.1093/cid/ciy776.

5. Caputo, R.; Asprea, M.; Giovannetti, L.; Messori, A. Nephrotoxicity of three formulations of amphotericin B: trial sequential analysis. Arch. Med. Sci. 2020, 16, 1493–1495, doi:10.5114/aoms.2020.93338.

6. Inselmann, G.; Inselmann, U.; Heidemann, H.T. Amphotericin B and liver function. Eur. J. Intern. Med. 2002, 13, 288–292, doi:10.1016/s0953-6205(02)00065-1.

7. Hui, X.; Baker, S.J.; Wester, R.C.; Barbadillo, S.; Cashmore, A.K.; Sanders, V.; Hold, K.M.; Akama, T.; Zhang, Y.K.; Plattner, J.J.;, et al. In vitro penetration of a novel oxaborole antifungal (AN2690) into the human nail plate. J. Pharm. Sci. 2007, 96, 2622–2631, doi:10.1002/jps.20901.

8. He, Z.; Huang, D.-C.; Guo, D.; Deng, F.; Sha, Q.; Zhang, M.-Z.; Zhang, W.-H.; Gu, Y.-C. Synthesis, fungicidal activity and molecular docking studies of tavaborole derivatives. Adv. Agrochem 2023, 2, 185–195, 10.1016/j.aac.2023.05.004.

9. Baker, S.J.; Zhang, Y.-K.; Akama, T.; Lau, A.; Zhou, H.; Hernandez, V.; Mao, W.; Alley; Sanders, V.; Plattner, J.J. Discovery of a new boron-containing antifungal agent, 5-fluoro-1,3-dihydro-1-hydroxy-2,1-benzoxaborole (AN2690), for the potential treatment of onychomycosis. J. Med. Chem. 2006, 49, 4447–4450, doi:DOI: 10.1021/jm0603724.

10. Adamczyk-Woźniak, A.; Borys, K.M.; Sporzyński, A. Recent Developments in the Chemistry and Biological Applications of Benzoxaboroles. Chem. Rev. 2015, 115, 5224–5247.

11. Lipner, S.R.; Scher, R.K. Onychomycosis: Treatment and prevention of recurrence. J. Am. Acad. Dermatol. 2019, 80, 853–867.

12. Nocentini, A.; Supuran, C.T.; Winum, J.-Y. Benzoxaborole compounds for therapeutic uses: a patent review (2010-2018). Expert Opin. Ther. Pat. 2018, 28, 493–504, doi:DOI: 10.1080/13543776.2018.1473379.

13. Rosen, T.; Gold, L.F.S. Antifungal drugs for onychomycosis: Efficacy, safety, and mechanisms of action. Semin. Cutan. Med. Surg. 2016, 35, S51–S55, doi:DOI: 10.12788/j.sder.2016.009.

14. Das, B.C.; Adil Shareef, M.; Das, S.; Nandwana, N.K.; Das, Y.; Saito, M.; Weiss, L.M. Boron-Containing heterocycles as promising pharmacological agents. Bioorg. Med. Chem. 2022, 63, 116748, 10.1016/j.bmc.2022.116748.

15. Dhawan, B.; Akhter, G.; Hamid, H.; Kesharwani, P.; Alam, M.S. Benzoxaboroles: New emerging and versatile scaffold with a plethora of pharmacological activities. J. Mol. Struct. 2022, 1252, 132057, 10.1016/j.molstruc.2021.132057.

16. Liu, S.; She, P.; Li, Z.; Li, Y.; Li, L.; Yang, Y.; Zhou, L.; Wu, Y. Drug synergy discovery of tavaborole and aminoglycosides against Escherichia coli using high throughput screening. AMB Express 2022, 12, 151, doi:10.1186/s13568-022-01488-6.

17. Adamczyk-Woźniak, A.; Cabaj, Małgorzata K. Dominiak, P.M.; Gajowiec, P.; Gierczyk, B.; Lipok, J.; Popenda, Ł.; Schroeder, G.; Tomecka, E.; Urbański, P.; Wieczorek, D.;, et al. The influence of fluorine position on the properties of fluorobenzoxaboroles. Bioorg. Chem. 2015, 60, 130– 135, doi:10.1016/j.bioorg.2015.05.004.

18. Borys, K.M.; Matuszewska, A.; Wieczorek, D.; Kopczyńska, K.; Lipok, J.; Madura, I.D.; Adamczyk-Woźniak, A. Synthesis and structural elucidation of novel antifungal N-(fluorophenyl)piperazinyl benzoxaboroles and their analogues. J. Mol. Struct. 2019, 1181, 587–598, 10.1016/j.molstruc.2019.01.018.

19. Borys, K.M.; Wieczorek, D.; Tarkowska, M.; Jankowska, A.; Lipok, J.; Adamczyk-Woźniak, A. Mechanochemical synthesis of antifungal bis(benzoxaboroles). RSC Adv. 2020, 10, 37187– 37193, doi:10.1039/D0RA07767D.

20. Wieczorek, D.; Kaczorowska, E.; Wiśniewska, M.; Madura, I.D.; Leśniak, M.; Lipok, J.; Adamczyk-Woźniak, A. Synthesis and Influence of 3-Amino Benzoxaboroles Structure on Their Activity against Candida albicans. Molecules 2020, 25, a5999, doi:10.3390/molecules25245999.

21. Adamczyk-Woźniak, A.; Tarkowska, M.; Lazar, Z.; Kaczorowska, E.; Madura, I.D.; Maria Dąbrowska, A.; Lipok, J.; Wieczorek, D.; Dąbrowska, A.M.; Lipok, J.;, et al. Synthesis, structure, properties and antimicrobial activity of para trifluoromethyl phenylboronic derivatives. Bioorg. Chem. 2022, 119, 105560, 10.1016/j.bioorg.2021.105560.

22. Adamczyk-Woźniak, A.; Komarovska-Porokhnyavets, O.; Misterkiewicz, B.; Novikov, V.P.; Sporzyński, A. Biological activity of selected boronic acids and their derivatives. Appl. Organomet. Chem. 2012, 26, 390–393, doi:10.1002/aoc.2880.

23. Wieczorek, D.; Lipok, J.; Borys, K.M.; Adamczyk-Woźniak, A.; Sporzyński, A. Investigation of fungicidal activity of 3-piperazine-bis(benzoxaborole) and its boronic acid analogue. Appl. Organomet. Chem. 2014, 28, 347–350, doi:10.1002/aoc.3132.

24. Athanasopoulos, A.; André, B.; Sophianopoulou, V.; Gournas, C. Fungal plasma membrane domains. FEMS Microbiol. Rev. 2019, 43, 642–673, doi:10.1093/femsre/fuz022.

25. Gizińska, M.; Staniszewska, A.; Kazek, M.; Koronkiewicz, M.; Kuryk, Ł.; Milner-Krawczyk, M.; Baran, J.; Borowiecki, P.; Staniszewska, M. Antifungal polybrominated proxyphylline derivative induces Candida albicans calcineurin stress response in Galleria mellonella. Bioorg. Med. Chem. Lett. 2020, 30, 127545, doi:10.1016/j.bmcl.2020.127545.

26. Adamczyk-Woźniak, A.; Borys, K.M.; Madura, I.D.; Michałek, S.; Pawełko, A. Straightforward synthesis and crystal structures of the 3-piperazine-bisbenzoxaboroles and their boronic acid analogs. Tetrahedron 2013, 69, 8936–8942, doi:DOI: 10.1016/j.tet.2013.07.102.

27. Madura, I.D.; Adamczyk-Woźniak, A.; Jakubczyk, M.; Sporzyński, A. 5-Fluoro-1,3-dihydro-2,1-benzoxaborol-1-ol. Acta Crystallogr. Sect. E Struct. Reports Online 2011, 67, o414–o415, doi:10.1107/S1600536811001632.

28. Reference method for broth dilution antifungal susceptibility testing of yeasts - approved standard.; Wayne, P., Ed.; Third edit.; Clinical and Laboratory Standards Institute, 2008;

29. The Gene Ontology resource: enriching a GOld mine. Nucleic Acids Res. 2021, 49, D325–D334, doi:10.1093/nar/gkaa1113.

30. Kolberg, L.; Raudvere, U.; Kuzmin, I.; Adler, P.; Vilo, J.; Peterson, H. g:Profiler-interoperable web service for functional enrichment analysis and gene identifier mapping (2023 update). Nucleic Acids Res. 2023, 51, W207–W212, doi:10.1093/nar/gkad347.

31. Zhao, H.; Palencia, A.; Seiradake, E.; Ghaemi, Z.; Cusack, S.; Luthey-Schulten, Z.; Martinis, S. Analysis of the Resistance Mechanism of a Benzoxaborole Inhibitor Reveals Insight into the Leucyl-tRNA Synthetase Editing Mechanism. ACS Chem. Biol. 2015, 10, 2277–2285, doi:10.1021/acschembio.5b00291.

32. Adamczyk-Woźniak, A.; Gozdalik, J.T.; Wieczorek, D.; Madura, I.D.; Kaczorowska, E.; Brzezińska, E.; Sporzyński, A.; Lipok, J. Synthesis, Properties and Antimicrobial Activity of 5-Trifluoromethyl-2-formylphenylboronic Acid. Molecules 2020, 25, a799, doi:10.3390/molecules25040799.

33. Kaczorowska, E.; Adamczyk-Woźniak, A.; Żukowska, G.Z.; Kostecka, P.; Sporzyński, A. Vibrational Properties of Benzoxaboroles and Their Interactions with Candida albicans’ LeuRS. Symmetry (Basel). 2021, 13, a1845, doi:doi.org/10.3390/sym13101845.

34. Arita, G.S.; Meneguello, J.E.; Sakita, K.M.; Faria, D.R.; Pilau, E.J.; Ghiraldi-Lopes, L.D.; Campanerut-Sá, P.A.Z.; Kioshima, É.S.; Bonfim-Mendonça, P. de S.; Svidzinski, T.I.E. Serial Systemic Candida albicans Infection Highlighted by Proteomics. Front. Cell. Infect. Microbiol. 2019, 9.

35. Soto, S.M. Role of efflux pumps in the antibiotic resistance of bacteria embedded in a biofilm. Virulence 2013, 4, 223–229, doi:10.4161/viru.23724.

36. Bataineh, M.T.A.L.; Soares, N.C.H.; Semreen, M.H.; Cacciatore, S.; Dash, N.R.; Hamad, M.; Mousa, M.K.; Salam, J.S.A.; Al Gharaibeh, M.F.; Zerbini, L.F.;, et al. Candida albicans PPG1, a serine/threonine phosphatase, plays a vital role in central carbon metabolisms under filament-inducing conditions: A multi-omics approach. PLoS One 2021, 16, 1–16.

37. Zhongle, L.; C., M.L. Candida albicans Swi/Snf and Mediator Complexes Differentially Regulate Mrr1-Induced MDR1 Expression and Fluconazole Resistance. Antimicrob. Agents Chemother. 2017, *61*, 10.1128/aac.01344-17, doi:10.1128/aac.01344-17.

38. Bandara, H.M.H.N.; Wood, D.L.A.; Vanwonterghem, I.; Hugenholtz, P.; Cheung, B.P.K.; Samaranayake, L.P. Fluconazole resistance in Candida albicans is induced by Pseudomonas aeruginosa quorum sensing. Sci. Rep. 2020, 10, 7769, doi:10.1038/s41598-020-64761-3.

39. Gomez, M.A.R.; Ibba, M. Aminoacyl-tRNA synthetases. RNA 2020, 26, 910–936.

40. Xu, H.; Fang, T.; Omran, R.P.; Whiteway, M.; Jiang, L. RNA sequencing reveals an additional Crz1-binding motif in promoters of its target genes in the human fungal pathogen Candida albicans. Cell Commun. Signal. 2020, 18, 1, doi:10.1186/s12964-019-0473-9.

41. McCall, A.D.; Kumar, R.; Edgerton, M. Candida albicans Sfl1/Sfl2 regulatory network drives the formation of pathogenic microcolonies. PloS Pathog. 2018, 1–28.

42. Fontaine, F.; Héquet, A.; Voisin-Chiret, A.-S.; Bouillon, A.; Lesnard, A.; Cresteil, T.; Jolivalt, C.; Rault, S. Boronic species as promising inhibitors of the Staphylococcus aureus NorA efflux pump: Study of 6-substituted pyridine-3-boronic acid derivatives. Eur. J. Med. Chem. 2015, 95, 185–198, 10.1016/j.ejmech.2015.02.056.

43. Durka, K.; Laudy, A.E.; Charzewski, Ł.; Urban, M.; Stępień, K.; Tyski, S.; Krzyśko, K.A.; Luliński, S. Antimicrobial and KPC/AmpC inhibitory activity of functionalized benzosiloxaboroles. Eur. J. Med. Chem. 2019, doi:10.1016/j.ejmech.2019.03.028.

44. Frisch, M.J.; Trucks, G.W.; Schlegel, H.B.; Scuseria, G.E.; Robb, M.A.; Cheeseman, J.R.; Montgomery, Jr., J.A.; Vreven, T.; Kudin, K.N.; Burant, J.C.;, et al. Gaussian 03, Revision C. 02 2004.

45. Lee, C.; Yang, W.; Parr, R.G. Development of the Colle-Salvetti correlation-energy formula into a functional of the electron density. Phys. Rev. B 1988, 37, 785–789, doi:10.1103/PhysRevB.37.785.

46. Wachters, A.J.H. Gaussian Basis Set for Molecular Wavefunctions Containing Third-Row Atoms. J. Chem. Phys. 1970, 52, 1033–1036, doi:10.1063/1.1673095.

47. Hay, P.J. Gaussian basis sets for molecular calculations. The representation of 3d orbitals in transition-metal atoms. J. Chem. Phys. 1977, 66, 4377–4384, doi:10.1063/1.433731.

48. Raghavachari, K.; Trucks, G.W. Highly correlated systems. Excitation energies of first row transition metals Sc–Cu. J. Chem. Phys. 1989, 91, 1062–1065, doi:10.1063/1.457230.

49. Binning Jr., R.C.; Curtiss, L.A. Compact contracted basis sets for third-row atoms: Ga–Kr. J. Comput. Chem. 1990, 11, 1206–1216, 10.1002/jcc.540111013.

50. McGrath, M.P.; Radom, L. Extension of Gaussian-1 (G1) theory to bromine-containing molecules. J. Chem. Phys. 1991, 94, 511–516, doi:10.1063/1.460367.

51. Curtiss, L.A.; McGrath, M.P.; Blaudeau, J.; Davis, N.E.; Binning, R.C.; Radom, L. Extension of Gaussian-2 theory to molecules containing third-row atoms Ga–Kr. J. Chem. Phys. 1995, 103, 6104–6113, doi:10.1063/1.470438.

52. McLean, A.D.; Chandler, G.S. Contracted Gaussian basis sets for molecular calculations. I. Second row atoms, Z=11–18. J. Chem. Phys. 1980, 72, 5639–5648, doi:10.1063/1.438980.

53. Krishnan, R.; Binkley, J.S.; Seeger, R.; Pople, J.A. Self-consistent molecular orbital methods. XX. A basis set for correlated wave functions. J. Chem. Phys. 1980, 72, 650–654, doi:10.1063/1.438955.

54. Staniszewska, M.; Zdrojewski, T.; Gizińska, M.; Rogalska, M.; Kuryk, Ł.; Kowalkowska, A.; Łukowska-Chojnacka, E. Tetrazole derivatives bearing benzodiazepine moiety-synthesis and action mode against virulence of Candida albicans. Eur. J. Med. Chem. 2022, 230, 114060, doi:10.1016/j.ejmech.2021.114060.

55. George, P.; Deborah, A.; Heather, K.; Abdulrahim, I. Time-Kill Assay and Etest Evaluation for Synergy with Polymyxin B and Fluconazole against Candida glabrata. Antimicrob. Agents Chemother. 2014, 58, 5795–5800, doi:10.1128/aac.03035-14.

56. Staniszewska, M.; Sobiepanek, A.; Gizińska, M.; Peña-Cabrera, E.; Arroyo-Córdoba, I.J.; Kazek, M.; Kuryk, Ł.; Wieczorek, M.; Koronkiewicz, M.; Kobiela, T.;, et al. Sulfone derivatives enter the cytoplasm of Candida albicans sessile cells. Eur. J. Med. Chem. 2020, 191, 112139, doi:10.1016/j.ejmech.2020.112139.

57. Łukowska-Chojnacka, E.; Kowalkowska, A.; Gizińska, M.; Koronkiewicz, M.; Staniszewska, M. Synthesis of tetrazole derivatives bearing pyrrolidine scaffold and evaluation of their antifungal activity against Candida albicans. Eur. J. Med. Chem. 2019, 164, 106–120, doi:10.1016/j.ejmech.2018.12.044.

58. Gómez-Gaviria, M.; Lozoya-Pérez, N.E.; Staniszewska, M.; Franco, B.; Niño-Vega, G.A.; Mora-Montes, H.M. Loss of Kex2 Affects the Candida albicans Cell Wall and Interaction with Innate Immune Cells. J. Fungi 2020, 6.

59. OECD Guidelines for the Testing of Chemicals, Section 2, Test No. 236: Fish Embryo Acute Toxicity (FET) Test;

60. Fiołka, M.J.; Mieszawska, S.; Czaplewska, P.; Szymańska, A.; Stępnik, K.; Sofińska-Chmiel, W.; Buchwald, T.; Lewtak, K. Candida albicans cell wall as a target of action for the protein– carbohydrate fraction from coelomic fluid of Dendrobaena veneta. Sci. Rep. 2020, 10, 16352, doi:10.1038/s41598-020-73044-w.

61. Martin, M. CUTADAPT removes adapter sequences from high-throughput sequencing reads. EMBnet.journal 2011, 17, doi:10.14806/ej.17.1.200.

62. http://www.candidagenome.org/).

63. Love, M.I.; Huber, W.; Anders, S. Moderated estimation of fold change and dispersion for RNA-seq data with DESeq2. Genome Biol. 2014, 15, 550, doi:10.1186/s13059-014-0550-8.

64. Carlson, M. Bioconductor/GO.db: A set of annotation maps describing the entire Gene Ontology 2019.

